# Can we infer mechanisms governing PMF generation and partitioning from existing experimental data? A model-based identifiability analysis

**DOI:** 10.64898/2026.11.23.767962

**Authors:** Josha Arthur Ebeling, Anna Matuszyńska

**Author notes:** Correspondence &.

## Abstract

Photosynthesis is a well-studied and essential biochemical process on Earth. It converts light energy into chemical energy by coupling the photosynthetic electron transport chain to the Calvin–Benson–Bassham (CBB) cycle. Central to this coupling is the proton motive force (PMF) across the thylakoid membrane. It drives ATP synthesis, and its partitioning into membrane potential (ΔΨ) and proton gradient (ΔpH) regulates photoprotection and electron flow. How processes in the thylakoid membrane shape the size and composition of the PMF is, however, difficult to measure directly, as the available signals, whether from gas exchange or spectroscopy, integrate several processes. Mechanistic models of PMF regulation have been used to simulate its behavior, but it has not been assessed how much information standard measurements contain about the underlying mechanisms. We extended a dynamic model of C3 photosynthesis to combine an explicit description of ΔΨ and ΔpH, thylakoid ion transporters and the CBB cycle, and fitted it jointly to non-photochemical quenching (NPQ), CO_2_ assimilation and the electrochromic shift measured on *Arabidopsis thaliana*. However, agreement between simulations and experimental measurements is not a sufficient condition for mechanistic inference. We therefore assessed which data are required to identify the model parameters. Progressive fitting showed that only the combination of all three datasets constrained the model sufficiently to reproduce all three measurements, and profile likelihood showed that all fitted parameters are constrained in at least one direction within their bounds. Sobol’ sensitivity analysis showed that the PMF is controlled mainly by the total chloride concentration and photosynthetic control in high light, which the data constrain, and by photosystem amounts and CBB enzyme capacities in low light, which they do not. As the functional amount of photosystems depends on thylakoid architecture, this points to dynamic structural measurements as a valuable additional constraint. Existing measurements can thus reveal the mechanisms that shape PMF dynamics, provided they are combined, but inferring PMF magnitude also requires knowledge of the photosynthetic machinery that sets its capacity.

## Introduction

Photosynthesis converts light energy into chemical energy, providing the carbon compounds that sustain plant growth and, ultimately, terrestrial food production. At the biochemical level, oxygenic photosynthesis in plants involves two closely coupled sets of reactions, namely the photosynthetic electron transport chain (PETC) and the Calvin-Benson-Bassham (CBB) cycle [1]. The two pathways are coupled through a shared pool of ATP and NADPH: the CBB consumes ATP and NADPH, and the PETC replenishes them, creating a complex supply-and-demand relationship [2, 3, 4]. The link between supply and demand is the proton-motive force (PMF) that forms across the thylakoid membrane, generated by the PETC under illumination. It consists of an electrical gradient (the membrane potential, ΔΨ) and a chemical gradient (the difference in luminal and stromal pH, ΔpH). Light energy captured by the antennae drives charge separation and, consequently, electron transport through the components of the PETC to NADP^+^, producing NADPH (linear electron flow, LEF) [5, 6] (see Fig. 1 **B**), thereby establishing the pH gradient across the thylakoid membrane. In addition, charge separation in both photosystems (PSII and PSI), electrogenic electron transfer in cytochrome b_6_f (cyt b_6_f), and charge translocation across the membrane via membrane transport proteins form the membrane potential component of the PMF[7, 8, 9]. While the PMF directly drives ATP synthesis via the ATP synthase [10], [11], it indirectly lowers NADPH production by slowing plastoquinol (PQH_2_) oxidation at cyt b_6_f (referred to as photosynthetic control) [12] LEF alone produces too little ATP relative to NADPH to meet the demand of the CBB cycle [13]. Several processes are assumed to balance this deficit, including cyclic electron flow (CEF) around PSI [14] and the water-water cycle [15]. CEF via the chloroplast NADH dehydrogenase-like (NDH) complex additionally pumps protons, generating more PMF per electron [16, 17]. How the PMF is partitioned between ΔΨ and ΔpH matters because the two components have different regulatory effects. Whilst ΔpH activates photoprotection and photosynthetic control, a large ΔΨ promotes charge recombination in PSII and singlet oxygen formation [18]. Importantly, both components are thermodynamically equivalent for ATP synthesis. Upon illumination, ΔpH builds up slowly over tens of seconds as the lumen buffer is titrated, whereas ΔΨ forms within milliseconds and is then partially dissipated through counterion movements across the membrane [19, 20]. Known counterions with described membrane transport proteins located on the thylakoid membrane are potassium (K^+^) and chloride (Cl^−^) [21]. VCCN1 is a voltage-gated chloride channel that opens in response to ΔΨ build-up, allowing Cl^−^ influx into the lumen [22]. ClCe is a CLC-family protein that also mediates Cl^−^ transport, though its precise function is debated [23, 24]. KEA3 is a K^+^/H^+^ antiporter that moves protons from the lumen to the stroma in exchange for K^+^. Because it is electroneutral, it lowers ΔpH without directly changing ΔΨ. Its activity is regulated by stromal pH and by the chloroplast energy status through its C-terminal domain [25, 26]. It is particularly relevant for short-term light acclimation, as it accelerates NPQ relaxation after transitions from high to low light [25]. A thylakoid K^+^ channel (TPK3) has also been proposed [27]. However, it was later proven to localize to the tonoplast [28]. Further, yet-unidentified transporters are likely to contribute.

**Figure 1.**
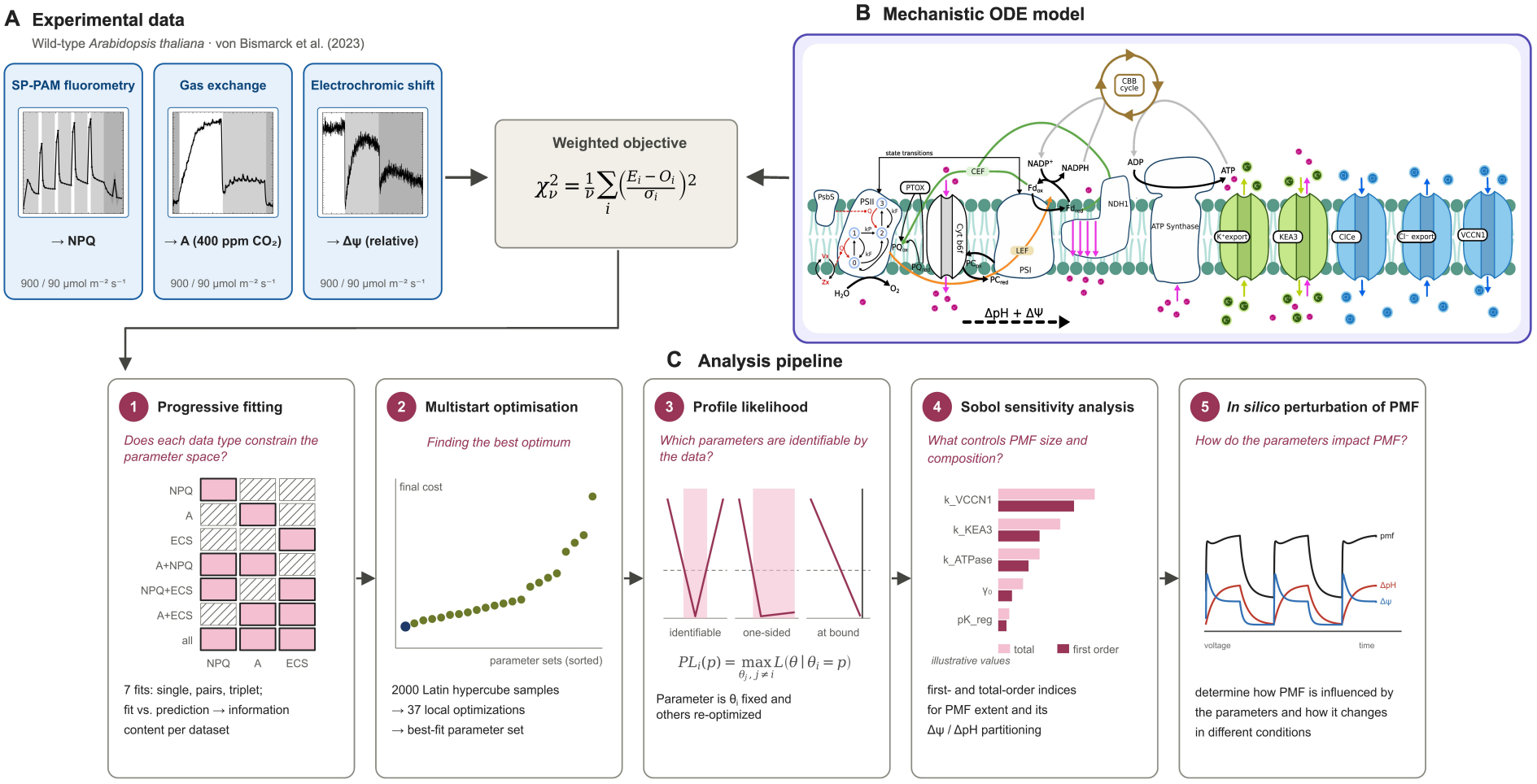
Workflow for inferring PMF regulation from combined photosynthesis measurements. **A** highlights the experimental data and the corresponding protocol of SP-PAM fluorometry, gas exchange, and electrochromic shift. The protocols are described in detail in the following section and are shown in Supplementary Figure S1. The schematic shown in **B** displays all the reactions and mechanisms our model includes: Photosynthetic electron transport chain, including linear electron flow and two cyclic electron flow pathways. At PSI, the Mehler pathway can convert excess energy into ROS. The PETC generates proton motive force, an electrochemical gradient, which drives ATP synthase. Five membrane transport proteins for ions, including VCCN1, ClCe, and a K^+^/H^+^ antiporter called KEA3, regulate chloride and potassium fluxes across the thylakoid membrane. ATP and NADPH are synthesized in the PETC and used in the Calvin-Benson-Bassham cycle, where Rubisco fixes CO_2_ onto RuBP. From there, the cycle produces sugar phosphates and regenerates RuBP. **C** represents the workflow of the study: First, we combine experimental and theoretical approaches by progressively fitting the model to the data. Next, we determine the parameter set that best represents the data by using multistart optimization. Once we find an optimum, we investigate parameter identifiability and their impact on PMF through profile likelihood and Sobol’ sensitivity analysis. To further complement Sobol’ analysis, we show how parameters with large total order Sobol’ indices influence PMF and its composition.

Despite this mechanistic knowledge, the size of each PMF component *in vivo* remains debatable. Estimates from electrochromic-shift (ECS) measurements suggest a substantial steady-state ΔΨ [19, 12], whereas other studies argue that overlapping absorbance changes distort these estimates and that ΔpH dominates the PMF [29, 30]. In high light, absorbed energy exceeds the CBB cycle capacity [31, 32]. The resulting over-reduction of the PETC can produce reactive oxygen species (ROS), which damage the photosystems. The main photoprotective mechanism is non-photochemical quenching (NPQ) in the PSII antenna, which dissipates excess energy as heat [33]. Its rapidly reversible component (qE) is triggered by lumen acidification through protonation of PsbS and activation of violaxanthin de-epoxidase in the xanthophyll cycle [34]. This makes NPQ, like photosynthetic control, a direct readout of ΔpH. Yet, measuring lumen pH *in vivo* remains challenging. Thylakoid-targeted pHluorin [35] has demonstrated light-dependent lumen acidification in *Arabidopsis thaliana*. However, transgene silencing limits its broader application [36, 37].

Because the PMF cannot be measured directly in leaves, its generation and partitioning have to be inferred from indirect observables. Three techniques are widely used to probe photosynthetic dynamics: Pulse Amplitude-Modulated (PAM) fluorometry, gas exchange measurements, and ECS. In PAM fluorometry, saturating light pulses transiently close PSII reaction centers. The resulting maximal fluorescence yields are used to derive NPQ and PSII operating efficiency (ΦPSII) [38]. Gas-exchange measurements record the difference in CO_2_ concentration between air entering and leaving the leaf chamber, from which net CO_2_ assimilation (A) is derived. Finally, the ECS is the change in carotenoid absorbance around 515 nm, which is proportional to the transmembrane electric field [39, 40, 41]. Its relaxation during short dark intervals is used to estimate the total PMF, its partitioning into ΔΨ and ΔpH, and the proton conductivity of the ATP synthase (gH^+^) [42, 19]. Each of these signals integrates several processes: net CO_2_ uptake combines carboxylation, photorespiration, and respiration [43]. Fluorescence yield reflects both photochemistry and quenching [44]. And the ECS reports ΔΨ, from which PMF partitioning can only be inferred from dark-interval relaxation kinetics, assuming that the absorbance change is exclusively electrochromic, an assumption that has been questioned [29, 30].

Mechanistic models provide a mathematical framework for investigating the contribution of individual processes to measured signals. Numerical simulations allow these contributions to be explored under systematically var-ied conditions, while statistical methods quantify parameter uncertainty, assess identifiability, and evaluate the sensitivity of model outputs to parameter changes [45]. Identifiability is especially critical here, as an agreement between simulations and experimental measurements is a necessary but not sufficient condition for mechanistic inference. If the parameters are not identifiable, distinct parameter configurations may reproduce the data equally well, and the contributions of individual processes to the measured signal remain undetermined. This requires distinguishing between structural identifiability, which concerns whether parameters can be uniquely determined in principle, and practical identifiability, which concerns whether the available data constrain individual parameters to finite and sufficiently narrow confidence intervals within their predefined bounds.

Most C3 photosynthesis models are tailored to the application for which they were developed, parameterized, and validated. Consequently, different models capture different aspects of photosynthetic regulation, with complementary features distributed across the literature. Steady-state models that describe the PETC and CBB cycle in detail, such as that of Johnson and Berry [46], enable quantitative interpretation of PAM fluorometry and gas-exchange measurements at steady state, but by construction cannot resolve the transient PMF dynamics captured by measurements under fluctuating light. Among dynamic models, worth citing are the model by Davis *et al*. [21] that includes ΔΨ and ion fluxes to study how PMF partitioning affects photoprotection and photodamage, and Li *et al*. [47], which describes the membrane potential, its regulation by specific transport proteins and the effects of their knockout. Both describe PMF dynamics in detail but simplify the CBB cycle and thus the demand side of photosynthesis. Morales *et al*. [48] developed a dynamic model of the PETC that includes PMF regulation, alternative electron transport, and Rubisco activation, focusing on the coupling between electron transport and carbon metabolism. They showed that it agrees with gas exchange, chlorophyll fluorescence, and absorbance measurements compiled from several studies, but it represents the PMF without explicit counterion fluxes through thylakoid transporters. Saadat *et al*. [49] linked the PETC to a full CBB description, thioredoxin regulation, and the Mehler reaction, but represented the PMF only through ΔpH. Addressing PMF generation and partitioning therefore requires a model that combines an explicit ΔΨ and ΔpH, specific thylakoid ion transporters, a mechanistic description of NPQ and a full CBB cycle, features that are so far distributed across these two groups of models. Moreover, all the above-mentioned models have been used in the forward direction to simulate the behavior of the PMF with parameters taken from the literature or fitted to individual datasets. The inverse question has not been addressed: how much information the standard measurements contain about the mechanisms of PMF regulation, and which of the underlying parameters they determine (1*C*). Although parameter identifiability has been examined for steady-state models fitted to gas-exchange data [50], to our knowledge, it has not been performed for dynamic models of PMF regulation.

Here we address this question by combining a dynamic model with progressive fitting and practical identifiability analysis, using published measurements on wild-type *Arabidopsis thaliana* [51]. We extend the C3 model of photosynthesis [49] to include a full PMF description, four ion channels and one antiporter, a dynamic photosystem description, and a dynamic description of both lumen and stroma protonation (Fig.1**B**). To avoid confounding the analysis with differences between plants, growth conditions, and protocols, all three measurements were taken from this single study, so our conclusions refer to one genotype, growth condition, and CO_2_ concentration. We then used it (i) to determine how strongly each measurement constrains the model and whether all three are needed; (ii) to assess the practical identifiability of its 32 fitted parameters; (iii) to quantify, by sensitivity analysis, which processes control the size and composition of the PMF under different light conditions.

## Methods

### 2.1. Mathematical model of C3 photosynthesis

We extended a previously published dynamic, kinetic model of C3 photosynthesis that links the PETC to a detailed description of the CBB cycle, including its regulation by thioredoxin [49]. We tested the performance of that model version against the measurements used in this study, to highlight the need for our applied changes (Fig S2). The model is a system of 43 ordinary differential equations (ODEs). Its time-dependent input is the photosynthetic photon flux density (PPFD, µmol photons m^−2^ s^−1^), and the CO_2_ concentration available to Rubisco is set by a parameter, the intercellular CO_2_ concentration (0.01326 mM C_*i*_, dissolved in the stroma, corresponding to 400 ppm, according to Henry’s law). In total, we added, removed, or modified 46 parameters, including 24 new parameters, 19 reactions, and 13 state variables. The supplementary material contains the complete model description, including all ODEs, rate equations, parameter values, and initial values (Table S2, S3, S4, S5).

#### 2.1.1. Model extensions: proton-motive force, ion transport and ATP synthase regulation

We extended the model in the following steps to describe the PMF and processes that generate and consume it. All added or modified parameters are listed in Supplementary Table S1.

##### Photosynthetic control

We introduced the slowing of PQH_2_ oxidation at cytochrome b_6_f upon lumen acidification, following [21]. The cyt b_6_f rate is multiplied by:

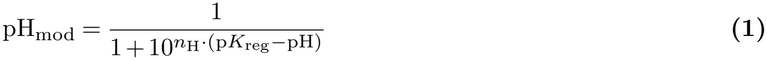

where *n*_H_ is a Hill coefficient and p*K*_reg_ the lumen pH of half-maximal inhibition. As luminal pH acidifies, pH_mod_ decreases and, in turn, slows PQH_2_ oxidation at cytochrome b_6_f. With photosynthetic control, the simulated lumen pH remained within the physiological range reported by [52].

##### Photosystem states

In the Saadat *et al*. (2021) model [49], the states of PSII and PSI were calculated under a quasi-steady-state assumption, which did not conserve the number of electrons transferred through the photosystems upon rapid transitions in light We now describe the photosystem states with explicit ODEs, after [53].

##### Luminal and stromal pH

The stromal pH, previously fixed at 7.8, is now a dynamic state variable, similar to luminal pH. Proton concentrations in each compartment are derived from the respective pH:

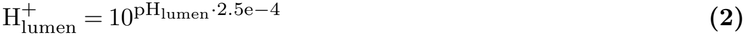

where 2.5e-4 represents the conversion factor between M and mmol/Mol Chl [54]. Consequently, the buffering constant *b*_H_, previously dimensionless, is now a buffer capacity: the amount of protons, in mmol (mol Chl)^−1^, required to lower the lumen pH by one unit. The same concept applies to stroma pH. We set the buffering capacity to 2.5 times the luminal buffering capacity so the stroma pH increases toward 8 upon illumination, as reported in Werdan *et al*. [55].

##### Membrane potential and ion transport

We introduced the membrane potential, ΔΨ, following Li *et al*. [47]. ΔΨ is proportional to the net charge translocated across the thylakoid membrane.

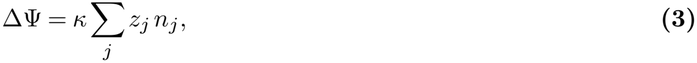

where *n*_*j*_ is the net amount of charged species *j* translocated into the lumen (mmol (mol Chl)^−1^), *z*_*j*_ its charge, and *κ* the potential generated per unit charge (V per mmol (mol Chl)^−1^). To allow counterion movement to downregulate ΔΨ, we included potassium and chloride and five thylakoid transport proteins (Fig. 1): the K^+^/H^+^ antiporter KEA3, the chloride channels VCCN1 and ClCe. Additionally, we assumed two hypothetical channels with unknown molecular identity to balance the one-directional potassium flux mediated by KEA3 and support ClCe in reinstating chloride homeostasis. The hypothetical potassium channel mediates K^+^ flux into the stroma at the onset of light, and the hypothetical chloride channel rebalances Cl^−^ between the compartments during relaxation in low light and darkness. The transporter rate equations are based on Li *et al*. [47], with modified regulation.

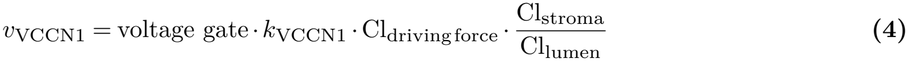

where Cl_drivingforce_ is the driving force of Cl^−^ across the thylakoid membrane.

Each transporter is regulated by a different variable: ΔΨ for VCCN1 and the hypothetical K^+^ channel, stromal ATP and stromal pH for KEA3, stromal ATP for ClCe, and the redox state of the PQ pool for the hypothetical Cl^−^ channel. Threshold-dependent activation is described by a sigmoid function:

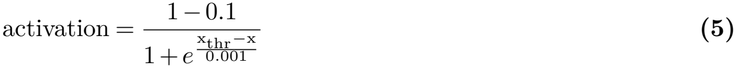

where *x* is the regulating variable, *x*_thr_ is its threshold value, and the numerator denotes the maximum activity (0.9). For the inhibitory regulators (e.g., stromal pH for KEA3), the sign of the exponent must be reversed.

##### Proton motive force

With the membrane potential ΔΨ (= *ψ*_lumen_*−ψ*_stroma_) and a dynamic ΔpH (= pH_stroma_ *−* pH_lumen_) in place, the free energy of transferring one mole of protons from the stroma to the lumen is given by the proton electrochemical potential difference

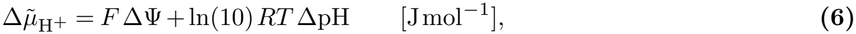

and the PMF is the same quantity expressed as an electrical potential,

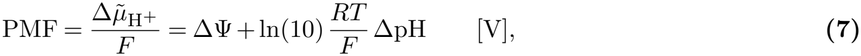

where ln(10)*RT /F ≈* 59 mV per pH unit at 25 ^°^C. In the equilibrium constants of the ATP synthase and cyt b_6_f reactions, the proton-gradient term Δ*G*_pH_ used in [49] was replaced by 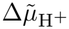 (Eq. 7), weighted by the number of protons translocated across the membrane (14*/*3 per ATP for the ATP synthase. And two per PQH_2_ oxidised for cyt b_6_f), so that both components of the PMF contribute to the thermodynamic driving force.

##### ATP synthase regulation

The ATP synthase is regulated in two ways. First, thiol modulation of the ATP synthase, i.e. the light-dependent reduction of a regulatory disulfide bridge in the *γ* subunit of CF_1_ by thioredoxin *f*, is approximated by a light-dependent switch acting on the active (reduced) fraction (*A*) of the enzyme:

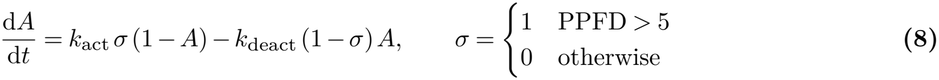

Second, the activity depends on the PMF, with half-maximal activation at a lower PMF for the reduced than for the oxidized enzyme [56]:

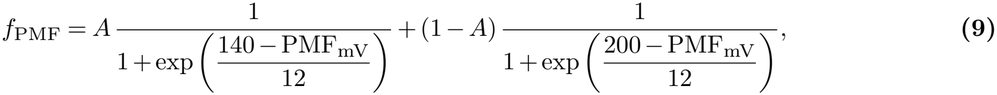

where PMF_mV_ is the PMF in mV. The half-activation values of 140 and 200 mV correspond to ΔpH values of 2.2 and 3.4, as reported by Junesch and Gräber (1987) [56]. This describes how much PMF is needed to activate the ATP synthase in the reduced and oxidized states. We increased the threshold from the reported 2.2 ΔpH.

##### Additional reactions

We added CEF via the chloroplast NADH dehydrogenase-like (NDH) complex [17]. Finally, we added an irreversible mass-action reaction that oxidizes thioredoxin, as a simplified representation of the Trx-like 2/2-Cys peroxiredoxin pathway, in which reduced thioredoxin is reoxidized independently of CBB cycle enzyme activation [57].

#### 2.1.2. Units

Units are consistent with previous versions of the model [49, 3], where we assume luminal concentrations are expressed in mmol (mol Chl)^−1^ and stromal concentrations in mM. We assume a chlorophyll content of 350 *·* 10^−6^ mol per m^2^ thylakoid membrane. The lumen volume we assume to be 0.0014 l m^−2^ and stroma 0.0112 l m^−2^ [54]. For species produced in one compartment and consumed in the other (ATP, NADPH, and H_2_O_2_), and for K^+^ and Cl^−^ transported across the thylakoid membrane, the stoichiometry includes a conversion factor that accounts for the compartment volumes.

### 2.2. Numerical simulation

We implemented the model in MxlPy v0.66.0 [58], a Python package for building and analyzing mechanistic models, and integrated it with RADAU [59]. Initial conditions are reported in the Supplement Table S5 The model is available in the mxlbricks package (mxlpy format), on mxlweb [60], which allows example simulations and simple parameter changes in a web browser without programming, and in the Greensloth database of photosynthesis models [61], from which it can be downloaded in several formats. The model and all code used for the analyses are available on GitHub.

### 2.3. Parametrization

We estimated parameters in two stages: global optimization with differential evolution, followed by local refinement of the best solution with a trust-region least-squares algorithm, both implemented in SciPy v1.18.1 [62]. For sampling, we used Latin hypercube sampling (LHS), which generates a near-random sample of parameter values from a multidimensional distribution. For multistart optimization, we drew 2000 parameter vectors within the parameter bounds by LHS (SciPy), selected 37 of them as starting points (see Results), and optimized each by least squares. Profile likelihoods were computed with pyPESTO v0.8.0 [63] and Sobol’ sensitivity indices with SALib v1.5.2 [64].

### 2.4. Experimental data

For all fitting activities reported here, we used published data from von Bismarck *et al*. (2023) [51]. The experiments were performed on *Arabidopsis thaliana*, grown in phytochambers under 12-h light at 90 µmol photons m^−2^ s^−1^ and 12-h darkness. We deliberately used published data from a single study, in which all three measurements were obtained from the same genotype grown under the same conditions and recorded with comparable light protocols. Combining measurements from different studies would introduce variation the model cannot represent, such as differences in growth light, plant age, temperature, instrumentation, and protocol. The model would compensate for these differences through its parameters, biasing both the parameter estimates and the identifiability analysis. Below, we briefly summarize the original experimental setup (1 *A*). Supplementary Figure S1 shows the light protocols, with the time windows used for fitting shaded.

#### 2.4.1. PAM fluorometry: NPQ

von Bismarck *et al*. (2023) [51] used the Maxi Imaging-PAM (Walz GmbH, Effel-trich, Germany) or an integrated diode emitter array spectrophotometer/fluorometer (IDEAspec, Hall *et al*. [65]). In PAM fluorometry, saturating light pulses (SPs) transiently close all PSII reaction centers, inducing maximal fluorescence yield [66]. From the measured yield values, quantities such as NPQ, the operating efficiency of PSII (Φ_PSII_), and the electron transport rate can be derived. NPQ was calculated as in Bilger *et al*. [67]:

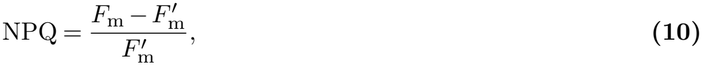

where *F*_m_ is the maximal fluorescence of the dark-adapted leaf and 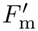 the maximal fluorescence during an SP in the light. After dark adaptation, the protocol starts with an SP (4000 µmol photons m^−2^ s^−1^) to determine *F*_m_, followed by low light (90 µmol photons m^−2^ s^−1^) with SPs after 80 s and then every 40 s. Four cycles follow, each consisting of 60 s of high light (900 µmol photons m^−2^ s^−1^) with SPs every 30 s and 240 s of low light. The protocol ends with 360 s of darkness with nine SPs.

#### 2.4.2. Gas-exchange: CO_2_ assimilation

Gas-exchange measurements were conducted on the youngest fully developed leaf with the Li-Cor 6400 gas exchange system (Li-Cor Biosciences, Lincoln, NE, USA; flow rate: 500 µmol s^−1^, CO_2_ concentration: 400 µmol mol^−1^, and humidity: 60%) coupled to the Maxi Imaging-PAM on whole plants, or the GFS-3000 open gas exchange system with LED array 3056-FL (Walz GmbH; flow: 600 µmol s^−1^, CO_2_ concentration: 400 µmol mol^−1^, humidity: 60%) [51]. This allows assessment of photosynthetic capacities such as carbon assimilation or stomatal conductance. In the model, net CO_2_ assimilation is calculated following the min-W approach from Lochoki and McGrath [68] based on Farquhar *et al*. [69]:

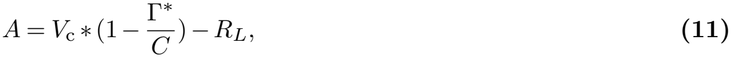

where *V*_c_ is the rate of Rubisco carboxylation. Γ^∗^ is the CO_2_ compensation point without non-photorespiratory CO_2_ release, *C* is the partial pressure of CO_2_ in proximity to Rubisco, and *R*_*L*_ is the rate of non-photorespiratory release of CO_2_, in the light. Gas exchange measurements were performed under atmospheric CO_2_ conditions of 400 ppm. The measurement protocol begins with a short dark adaptation period of 32 s, followed by 568 s of actinic light at 900 PPFD. It then transitions to 90 PPFD for 568 s, followed by 300 s of darkness.

#### 2.4.3. Electrochromic shift: membrane potential

The ECS is an absorbance increase at 515-520 nm in plant carotenoids. It was measured at six wavelengths (475, 488, 505, 520, 535, and 545 nm) with the IDEAspec. The signal was used to study proton flow through ATP synthase, and the decay rate was used to study qE, membrane conductivity, and ΔΨ [70]. For our analyses, we consider the derived ΔΨ dynamics, which is the deconvoluted ECS signal (ECS(ΔA, rel)). The measurement does not directly quantify the membrane potential in voltage units. Therefore, only the qualitative behavior would be comparable. For quantitative comparisons, we solve a system of linear equations for a factor “a” and an offset “b” that convert the model units to the measurement units. The ECS protocol consists of four iterations of the same three phases with a 90 PPFD phase before and after the four iterations. Each iteration consists of a high-light phase at 900 PPFD for 60 s, a 20 s low-light phase at 90 PPFD, and a 30 s darkness phase, which we simulate at 1 PPFD.

### 2.5. Objective function and goodness of fit

The three measurements differ in protocol length, number of data points, and numerical scale. To weight them comparably in our fitting routines, each residual was normalized by the measurement uncertainty and by the number of data points in its dataset,

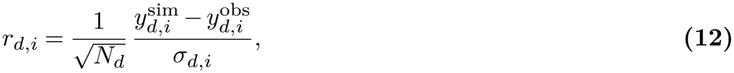

where 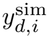 and 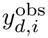 are the simulated and observed values of data point *i* in dataset *d, N*_*d*_ the number of data points in dataset *d*, and *σ*_*d,i*_ its uncertainty. The objective function minimized during fitting is 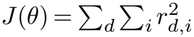. For NPQ and CO_2_ assimilation, *σ*_*d,i*_ is the standard error of the mean across replicates, smoothed with a centered moving average. For the ECS, we used only one replicate. We therefore estimated a constant noise level from the differences between consecutive data points, 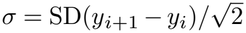, assuming uncorrelated noise between consecutive points.

The goodness of fit for each dataset was quantified by the reduced *χ*^2^,

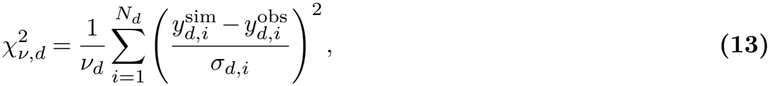

where *v*_*d*_ is the degrees of freedom and *v*_*d*_ = *N*_*d*_ *− p*, with *p* the number of fitted parameters, and *N*_*d*_ the number of data points. Because *σ*_*d,i*_ was estimated differently for each dataset, 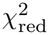 values are comparable only within a dataset, not between datasets.

#### 2.5.1. Selection of fitted parameters

Most model parameters were taken from the literature or from previous versions of the model. We fitted 32 parameters that fall into three groups: (i) parameters with a wide range of reported values, (ii) parameters that have not been measured, and (iii) phenomenological parameters without a direct physical counterpart, such as the factors *γ*_0_–*γ*_3_ that scale the four states of the NPQ quencher. Most of the parameters introduced in this model version belong to groups (ii) and (iii). Supplementary Table S1 lists the fitted parameters and their bounds.

#### 2.5.2. Profile likelihood

By performing profile likelihood at a previously determined optimum 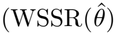, Eq. (12)), we investigate the identifiability of the fitted parameters at the optimum. For each parameter, the profile is obtained by fixing the parameter (*θ*_*i*_) slightly away from its previously determined optimum and reoptimizing the others, minimizing WSSR [71]:

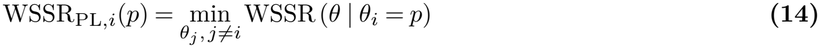

We then compare WSSR_PL,*i*_(*p*) to 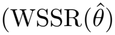 by taking their difference. Because WSSR has no meaningful scale, interpreting differences in WSSR is difficult. Therefore, we calculate s^2^ as an a posteriori estimate of the noise and use it to transform ΔWSSR to a noise scale, effectively telling us how much the fit changed in noise units.

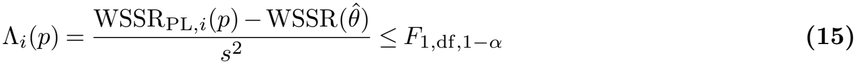

Because of the densely sampled time series of ECS and gas exchange, the residuals are not independent. Because of this high sampling density, compared to the low-density sampled NPQ, we keep weighting the residuals by the lengths of the corresponding datasets. With those two assumptions, we can only approximate a true F-distribution, including confidence intervals. However, the plots can still be interpreted in terms of practical identifiability, as Λ_*i*_(*p*) is still a valid measurement. To visualize the identifiability, we plot Λ_*i*_(*p*) against the perturbation of one parameter. The shape of the resulting curve indicates the parameter’s identifiability. A V- or U-shape indicates that the parameter is identifiable from the data. A rise on one side suggests non-identifiability, but different data might increase identifiability. A flat profile indicates either a structural non-identifiability, in which changes in the parameter are fully compensated by other parameters, or a negligible influence of the parameter on the fitted outputs.

#### 2.5.3. Sobol’ sensitivity analysis

Sobol’ sensitivity analysis is a global, variance-based sensitivity analysis applied to mathematical models to understand how variations in the parameter space affect the model output [72], [73]. We present total order indices, which describe how much of the output variance (Var(Y)) can be contributed to parameter *X*_*i*_, including all interactions. In the analysis, all parameters of interest are sequentially varied from their prior-determined optimum within a factor of two. Except for pH-related parameters, because varying them within a factor of two would span values from approximately three to twelve, which is far beyond physiological values in the lumen or stroma. Therefore, we vary these parameters in the range of *±* 0.5. The resulting change in model output is recorded (Var_*X i*_ (*Y* |*X*_~*i*_)). This variation is repeated for all possible parameter settings and averaged 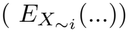. The average of the variance is then divided by the total variance (Var(Y)).

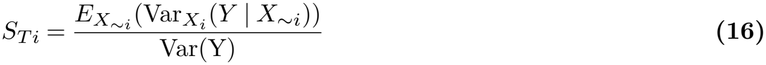

Such a global sensitivity analysis can be very useful in dissecting how the model output is determined, or rather, which parameters determine the output.

## Results

### 3.1. Progressive fitting

We first tested whether each measurement contributes distinct information by fitting the model progressively: first to each dataset alone, then to each pair, and finally to all three jointly. We define progressive fitting as a series of fits in which we consecutively increase the number of datasets the model is fitted to. Each fit started from 16 parameter sets selected from 1000 LHS (Methods), and every fit was then used to simulate all three measurements (Supplement Figure S3). Fig. 2 visualizes the quantitative assessment of each fit, showing the 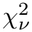 between the simulation and data for the measurement in each column. The lower the 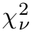, the better the agreement between the simulation with the fitted parameter set and the experiment. Fits to single datasets or pairs reproduced the fitted data but predicted the remaining measurements poorly. The NPQ-only fit generalized best (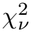 of 584.8 for assimilation and 68.7 for ECS), whereas the ECS-only fit predicted the other measurements worst. Adding a second dataset did not resolve this: fitting NPQ together with assimilation increased the NPQ’s and A’s 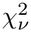 2-fold, and ECS prediction remained poor. Fitting NPQ with ECS left assimilation unpredicted 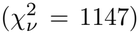. Only the joint fit reproduced all three measurements, indicating that each dataset constrains parameters that the others leave undetermined.

**Figure 2.**
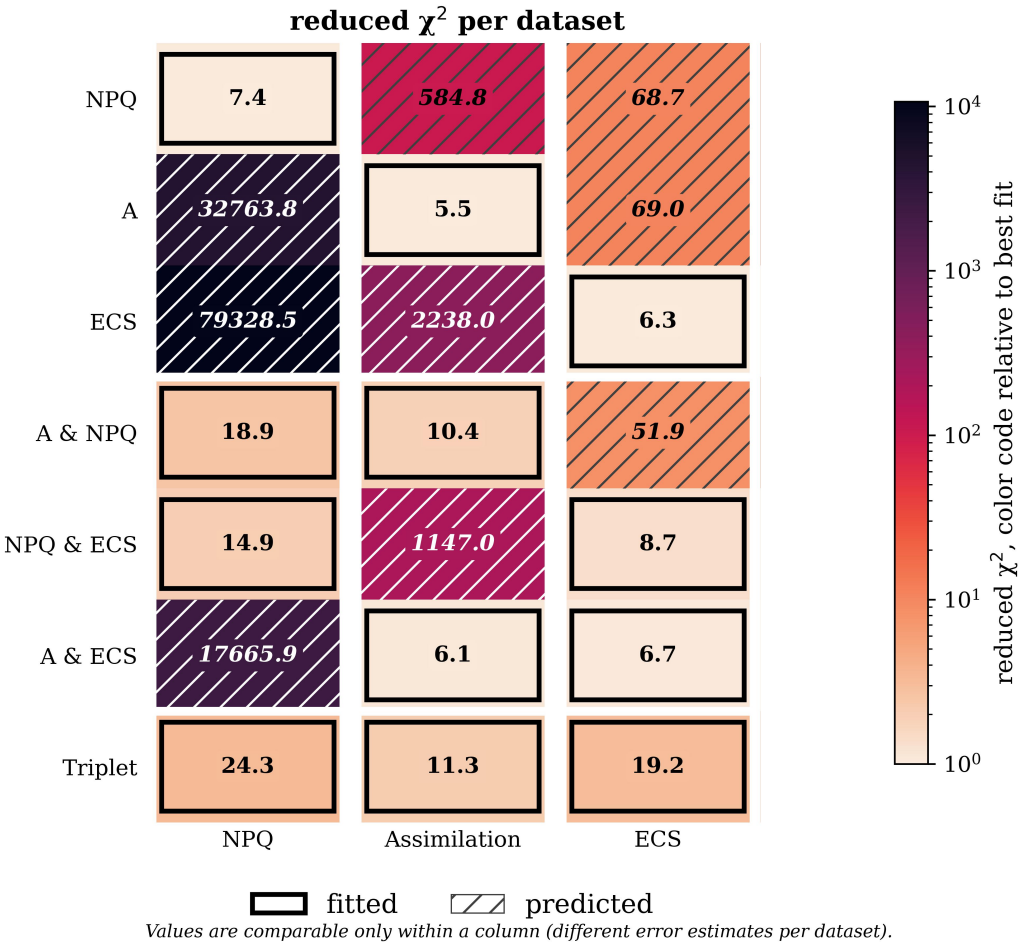
No single measurement or pair of measurements constrains the model sufficiently to predict the remaining ones. The model was fitted to each measurement alone (rows 1–3), to each pair (rows 4–6) and to all three jointly (row 7), and then used to simulate all three measurements. Each cell shows the reduced *χ*^2^ between simulation and data for one measurement (columns: NPQ, CO_2_ assimilation, ECS). Outlined cells are fitted data, hatched cells are predictions of data not used for fitting. Predictions of unseen data were consistently worse than fits, by up to three orders of magnitude, whereas the joint fit reproduced all three measurements with intermediate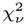.

The joint fit, however, did not reach the best 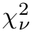 for any individual measurement, revealing a trade-off between datasets: parameter values that best explain one measurement are partly incompatible with the others. In a 32-dimensional parameter space with such competing objectives, the cost landscape is likely to contain several local optima. We therefore extended the search for the joint fit with a multistart optimization (Section 3.2).

### 3.2. Multistart optimization

We drew 2,000 parameter sets by LHS from the same parameter bounds as in progressive fitting and selected 37 of them as starting points: the 18 with the lowest cost before optimization, nine chosen iteratively to maximize the Euclidean distance to the nearest already selected set, nine chosen at random, and the joint fit from progressive fitting. Each starting point was optimized by least squares, and the resulting final costs were compared (Fig. 3).

**Figure 3.**
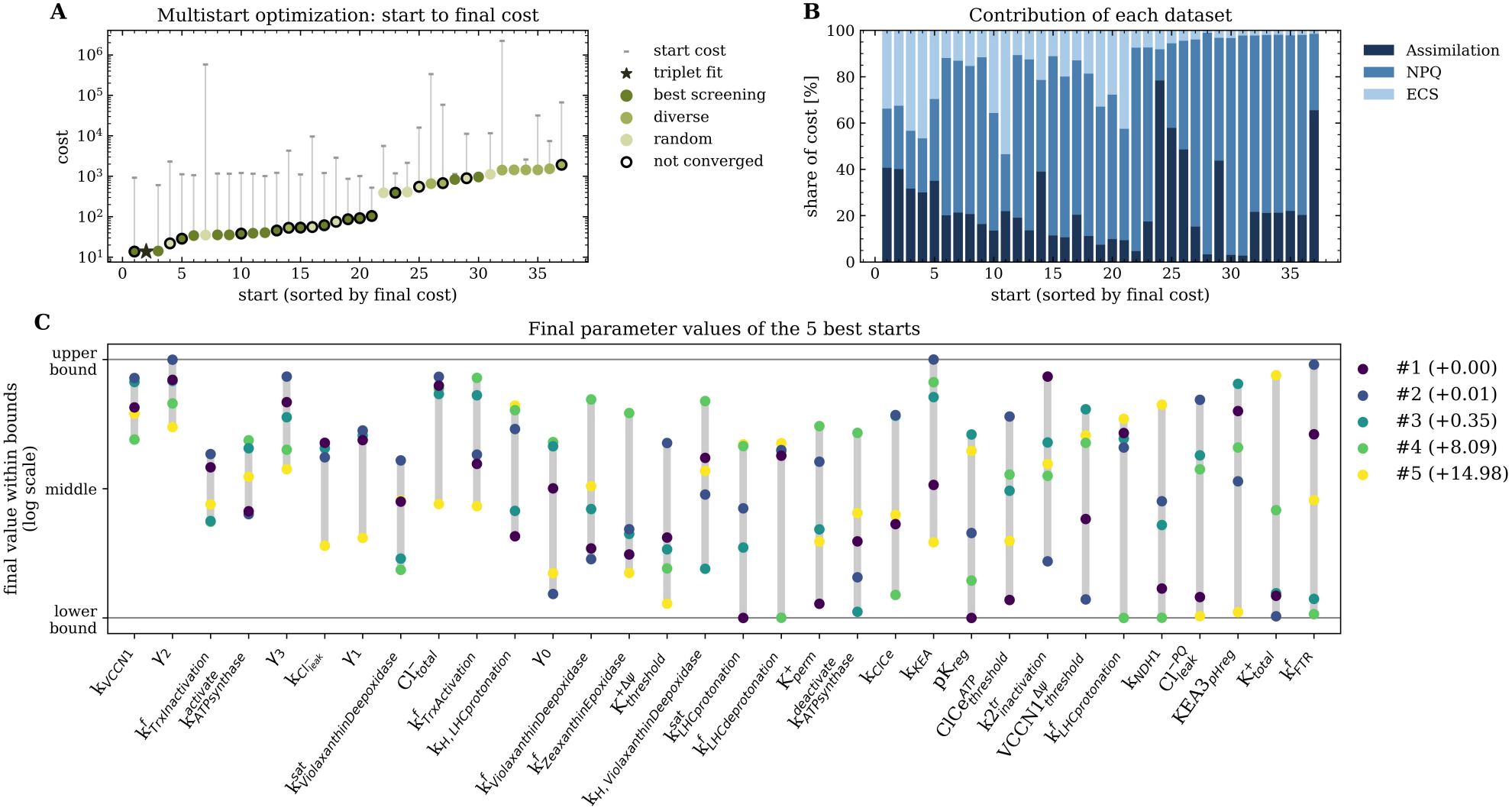
Multistart optimization reveals a plateau of near-equivalent optima with widely differing parameter values. **(A)** Final cost of the 37 starting points after least-squares optimization, sorted from lowest (left) to highest (right). Colors indicate how each starting point was selected from 2000 Latin hypercube samples: lowest initial cost (dark olive, *n* = 18), maximal Euclidean distance to already selected sets (olive, *n* = 9), and random (light olive, *n* = 9). The star marks the joint fit from progressive fitting. Grey lines connect each set to its cost before optimization. Black circles mark sets for which optimization stopped at the maximum number of iterations. Only the top three sets reached a final cost below 20. The lowest cost came from a set selected for its best initial performance. **(B)** Contribution of each dataset to the final cost of each set in **A**, in the same order (navy: assimilation; steel blue: NPQ; light blue: ECS). The shares of Assimilation and ECS increase toward lower costs. Whereas NPQ contributes the majority to most higher final costs. As final cost decreases, all three measurements contribute almost equally. In the four best sets, assimilation accounts for about 30-40%, ECS for 35-45% and NPQ for 25% of the cost. **(C)** Fitted parameter values of the five best sets, displayed within the parameter bounds on a logarithmic scale. Grey bars span the range across the five sets, and parameters are sorted from smallest to largest spread.

Final costs formed distinct clusters (Fig. 3A), with the worst-performing parameter sets originating from the random and diverse categories, scoring a final cost over 390. Eight sets ended at costs of 50–110 (five selected by initial cost, two by random and one by distance). The next ten parameter sets settled between 20–45 and comprised eight sets selected by initial cost and two randomly. Below are the top three performing sets, all with a cost below 15. The lowest final cost (13.84) was reached by a set selected by initial cost, for which optimization had not fully converged.

Figure 3B shows each dataset’s contribution to the final cost for each parameter set. In some of the worst-performing sets, NPQ accounts for most of the cost, reaching 95% in sets 31 and 32. For some sets, A takes over: In sets 24 and 25, it contributes 60-80% of the cost. ECS contributes the least up until set 21. Here, it contributes 40% and never falls below 10% for other sets again. As the final cost decreases, the shares of assimilation and ECS increase. In the five best sets, assimilation accounted for about 75%, ECS for 20%, and NPQ for 5% of the cost.

Despite similar costs, the three best sets differed in their parameter values (Fig. 3C). Parameters with the smallest spread, from *k*_VCCN1_ to *γ*_3_, converged to a similar region regardless of the starting point. Most parameters with a large spread formed two or more groups of values. For example, 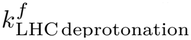, where all values except the fourth cluster between the middle and upper bounds. Other parameters, such as the forward rate constant of the Ferredoxin-Thioredoxin reductase 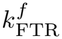, showed no grouping and spanned their full range.

In summary, the multistart optimization showed that few starting points converge to a plateau of similar costs, independent of how they were selected, and that the remaining cost in the best parametrizations can be equally attributed to all three measurements. At the same time, the three best parametrizations differ only marginally in cost but considerably in some parameter values, indicating that not all parameters are uniquely determined by the data. In the following, we therefore select the best-fitting parametrization and analyze its agreement with the data before assessing parameter identifiability in detail.

### 3.3. Reproduction of the experimental data with the best fit

We simulated all three measurement protocols using the parameter set yielding the lowest cost from the multistart optimization (Fig. 4). The model reproduced the magnitude and almost all dynamics of all three measurements: the NPQ induction and relaxation in each light cycle, the induction of CO_2_ assimilation and its transitions to low light and darkness, and the light-induced changes of the ECS signal.

**Figure 4.**
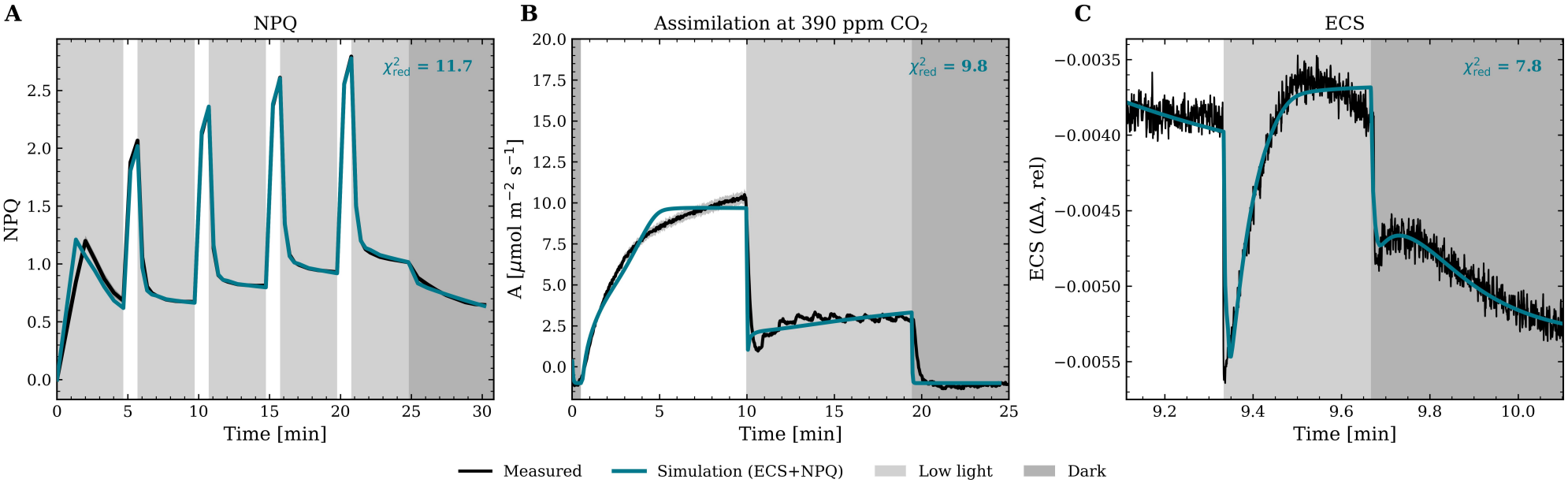
A single parametrization reproduces NPQ, CO_2_ assimilation and the ECS. Measured data (black) and simulations with the best parametrization from the multistart optimization (blue) for **A** NPQ, (B) net CO_2_ assimilation at 400 ppm CO_2_, and **C** the ECS-derived signal. Shading indicates low light (light grey) and darkness (dark grey); white areas are high light (900 PPFD).

The largest deviations occurred during NPQ 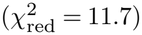 induction in the first low-light phase and in carbon assimilation 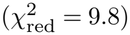 after minute five in the protocol. The model fails to slowly converge to steady-state assimilation and saturates too fast and too low.

### 3.4. Profile likelihood

A good agreement with data does not imply, though, that the parameters are determined by it. Before interpreting individual parameters mechanistically, we therefore quantified, for each fitted parameter, the range of values compatible with the data, using profile likelihood [45, 71]. We compare the performance of the original optimum and a newly found optimum when we perturb parameter *θ*_*i*_ and optimize all others. The perturbation is performed in both directions away from the original parameter value. We plot the profile likelihood, Λ_*i*_(p), against the perturbation, as in Figure 5, for all 32 fitted parameters. Because of the high frequency at which carbon assimilation and ECS were sampled, the residuals are not independent. Therefore, the critical value and confidence intervals are only approximate. The shape of the resulting profile still indicates identifiability: a V- or U-shaped profile that changes Λ_*i*_(p) drastically and crosses the approximate threshold on both sides indicates a well-identified parameter, a profile that crosses it on one side only indicates a parameter constrained in one direction, and a flat profile indicates a non-identifiable parameter.

**Figure 5.**
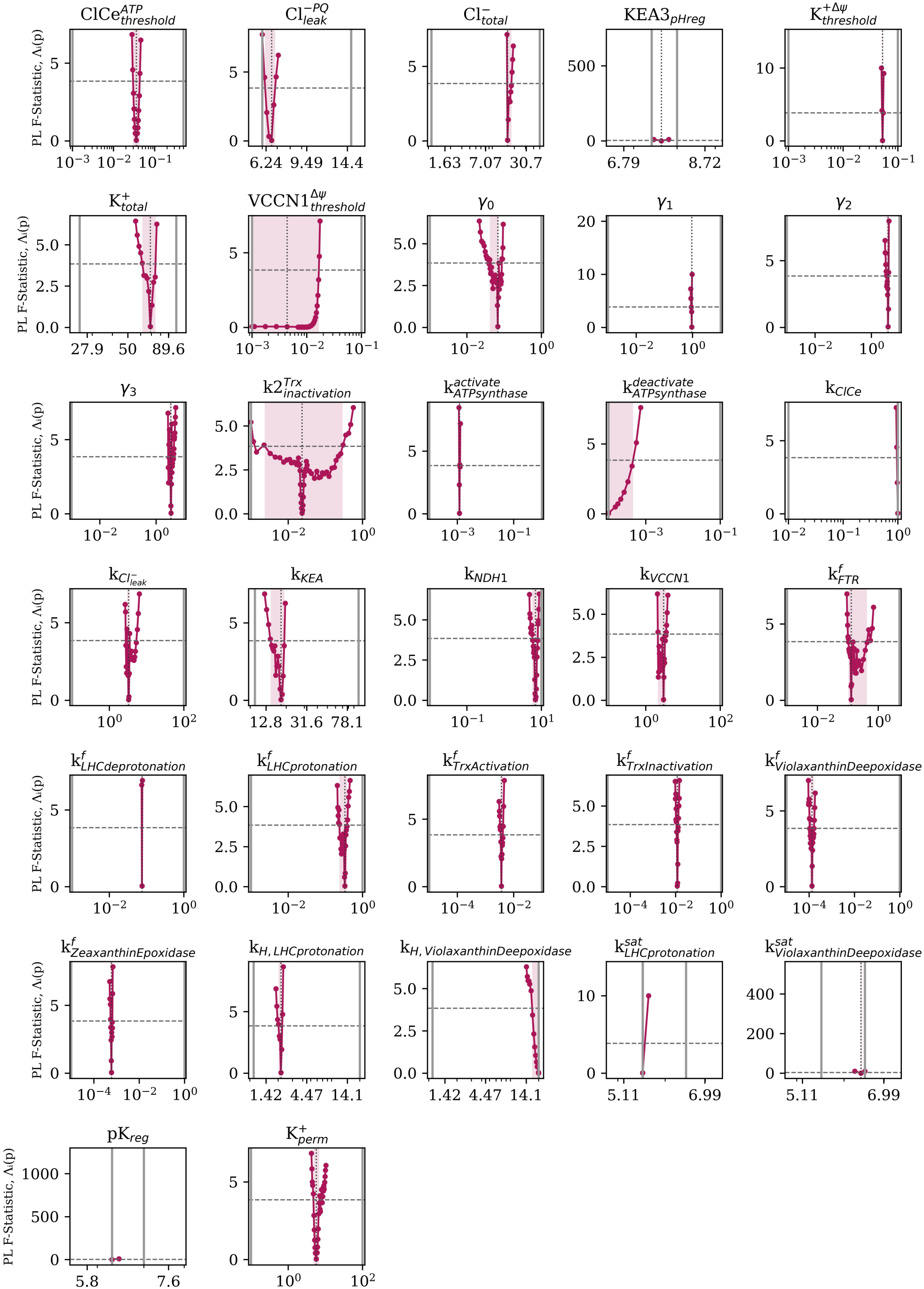
All fitted parameters are constrained by the combined data in at least one direction. Profile likelihoods of the 32 fitted parameters. Each parameter was fixed at values around its optimum (x-axis, logarithmic scale) while all others were reoptimized. The y-axis shows the difference Λ_*i*_(p) between those optimizations and the original optimum. Grey solid lines mark the parameter bounds, the dotted line the optimum, and the dashed line the approximate threshold of 3.84. Magenta areas indicate the confidence intervals. Note: Narrow profiles appear as near-vertical lines because the x-axis spans several orders of magnitude.

The dashed grey horizontal line is the chosen approximate critical value of 3.84, which marks the 95th percentile of the F distribution with one degree of freedom and a second with > 200. Using this critical value, we can determine the confidence intervals (magenta shaded areas), which mark the parameter range that reproduces data statistically indistinguishable from the best optimum. If Λ_*i*_(p), however, crosses the threshold, the fit deteriorates beyond the approximate 95% confidence level. Within the chosen parameter bounds, none of the 32 fitted parameters had a flat profile: every parameter was constrained by the data in at least one direction (Fig. 5). For 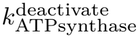, *k*_ClCe_, 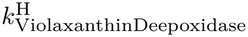,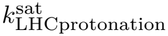, *pK*_reg_ optimal parameter value settled at the pre-defined bound. 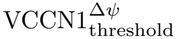 is the only parameter that exhibits unidentifiability on one side of the optimum, while the other side is well defined. As long as 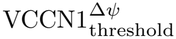 falls below the right confidence interval, it reproduces the data as well as the optimum.

### 3.5. Sobol’ sensitivity analysis

Profile likelihood shows how well the data determine each parameter, but not how strongly each parameter affects the size and composition of the PMF, which cannot be measured directly. We therefore performed a global Sobol’ sensitivity analysis [73, 72], which decomposes the variance of a model output into contributions of individual parameters and their interactions. The total-order index *S*_*T,i*_ is the fraction of output variance attributable to parameter *θ*_*i*_, including all its interactions with other parameters. In addition to the fitted parameters, we perturb PSII and PSI amounts, rate constants of CBB enzymes, cyclic electron flow, and ATP synthase. From the fitted parameters, we exclude several scaling parameters that would not correspond to identifiable biological processes (e.g., the four-state quencher parameters). Figure 6 shows the total-order indices for PMF across the three measurement protocols.

**Figure 6.**
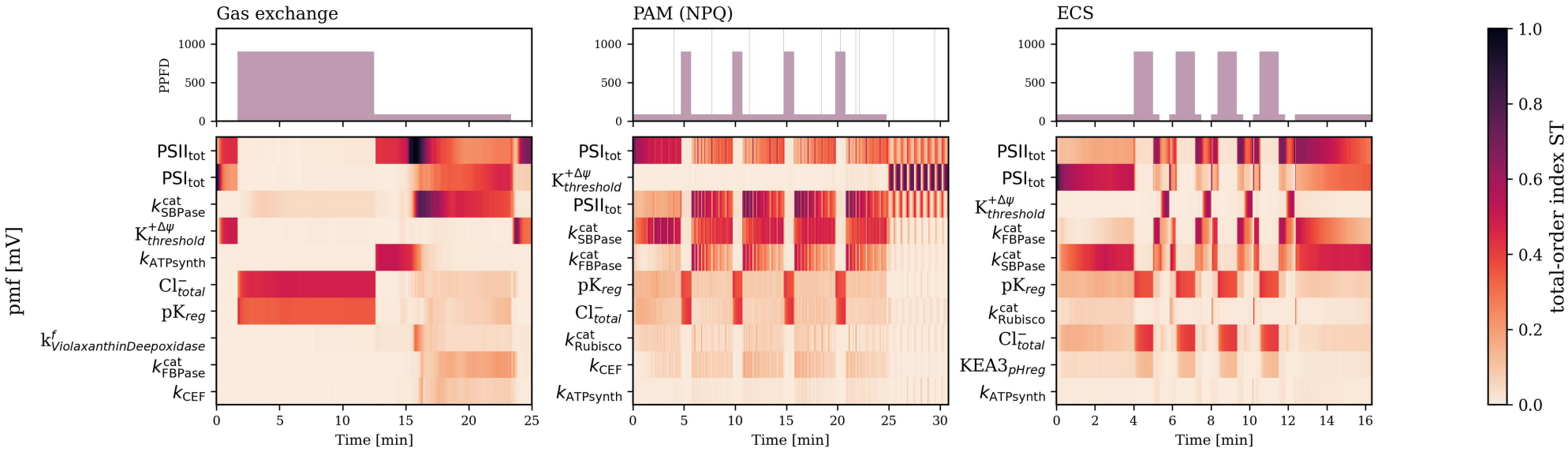
The PMF variability can be mostly accounted for by ion capacity and photosynthetic control in high light and in low light by photosystem amounts and CBB cycle capacity. Time-resolved total-order Sobol’ indices (*S*_*T*_) of the PMF for the gas-exchange, NPQ, and ECS protocols (light protocols shown above). Rows show the 10 parameters with the highest indices in each protocol. Fitted and non-fitted parameters were varied within a factor of 2 of their values, except pH-like parameters, which were varied within ± 0.5. Indices are less reliable where the PMF and its variance approach zero, e.g. in darkness.

Across the three protocols, total chloride concentration and *pK*_reg_ are the primary sources of PMF variance under high light. Under low light, however, the amounts of both Photosystems and catalytic rate constants of CBB enzymes, such as FBPase and SBPase, dominate PMF variability. By performing a similar set of parameter perturbations and simulations as in the Sobol’ analysis, we investigate how those parameters and their interactions shape PMF generation and composition.

### 3.6. Predicted PMF partitioning

Finally, using the best parametrization, we predict how the PMF partitions between ΔΨ and ΔpH, which none of the three measurements report directly (Fig. 7). In addition, we show how the six parameters with the highest average total-order indices in the ECS protocol influence PMF dynamics. Each parameter is perturbed within the same bounds as in the Sobol’ analysis. To align with the total-order indices, we simulated ECS for each parameter variation across 200 parameter combinations and averaged over those combinations to reflect the parameter’s impact on output variance and its interactions.

**Figure 7.**
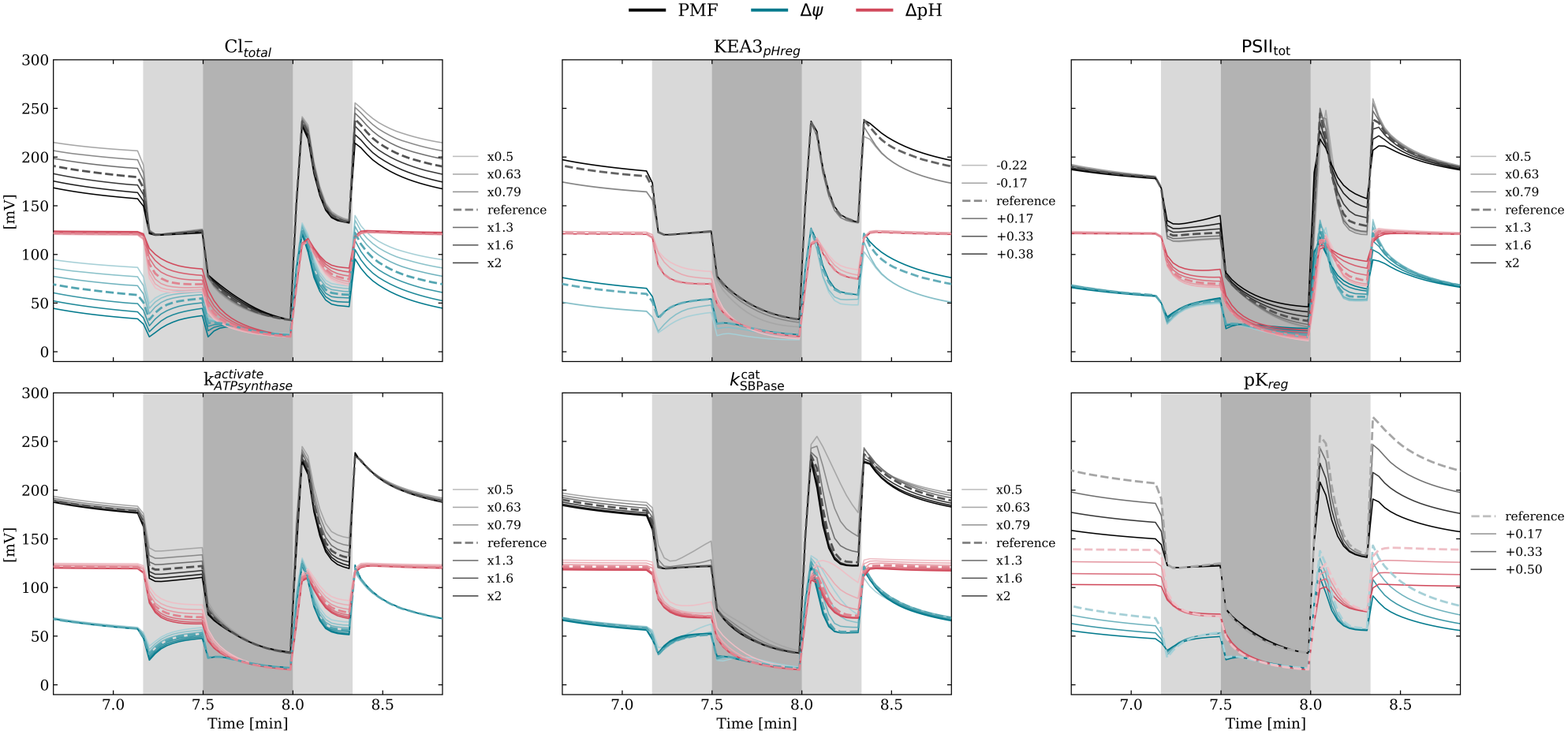
Predicted partitioning of the PMF into ΔΨ and ΔpH. PMF (black) decomposed into ΔΨ (blue) and ΔpH (red), during one cycle of the ECS protocol for the best parametrization and for variations of the six parameters with the highest total-order Sobol’ indices. Each variation was averaged over 200 combinations of the other parameters. White: high light (900 PPFD); light grey: low light (90 PPFD); dark grey: darkness.

Generally, in high light, ΔpH dominated the PMF, accounting for about 60% at steady state. In low light, ΔΨ and ΔpH contributed almost equally. At light transitions, the partitioning changed transiently: upon a decrease in light, ΔΨ collapsed within seconds while ΔpH relaxed more slowly. At an increase in light, ΔΨ rises higher than ΔpH and dissipates quickly, whereas ΔpH is less downregulated and, especially in high light, does not collapse at all.

Across the different parameters, we see different impacts on PMF and its composition. The total chloride ion concentration impacts both the membrane potential and the pH gradient. However, the membrane potential is affected under all light conditions, whereas the pH varies most when transitioning to lower light intensities. In high light, there is no difference. PSII’s total concentration causes PMF dynamics to vary in transients, showing a similar effect on ΔpH and ΔΨ. KEA3_pHreg_ and 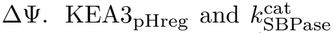 influence ΔΨ and ΔpH only when their value is decreased. They both increase ΔpH, however, only 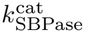 increases ΔΨ as well. The ΔΨ and ΔpH are both elevated during low-light conditions when 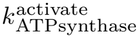 is varied.*pK*_reg_, the *pK*_a_ value for photosynthetic control, mainly alters the steady state of ΔΨ and ΔpH in light, although it also changes the dark-to-low-light transient.

## Discussion

Can we infer mechanisms governing PMF generation and partitioning from existing experimental data? Whereas previous models have simulated PMF regulation with assumed parameters, our results show which of these mechanisms the data actually constrain: no single measurement studied here constrained the model sufficiently, whereas jointly they constrained every fitted parameter in at least one direction, including those of thylakoid ion transport. At the same time, the magnitude of the PMF was controlled mainly by photosynthetic control and ion concentrations (constrained by the measurements) but also by photosystem amounts and CBB cycle enzymes, parameters that these measurements did not constrain.

As the three measurements we used quantify distinct parts of photosynthesis, each measurement constrains the model differently. Figure 2 highlights this: predictions on unseen data always performed at least twice as poorly as when the model is fitted to that data. Profile likelihood showed further that the data contain information about every parameter used in the fit, as no parameter exhibited a completely flat profile within its bounds. From this, we can draw conclusions about several parameters and their meaning, starting with the ion transport proteins. VCCN1 is well characterized in the literature and reported to be voltage-gated. The data indicate that the voltage gate should lie below 20 mV, relative to the model’s charge-to-voltage conversion, and the kinetic constant is 9.5 s^−1^. ClCe and the unknown chloride channel both fulfill a similar role. Therefore, we cannot conclude which channel is responsible, as excluding one could be compensated by the other when reoptimizing. We can only say that a chloride flux with a kinetic constant of the magnitude of both channels is needed to reproduce the data. For potassium, KEA3 and an unknown voltage-gated potassium channel are implemented in the model. Both parameters of the unknown channel are well identified by the data, suggesting a potassium flux of that order of magnitude, opening slightly delayed compared to VCCN1 at 52 mV and with a rate constant of 5.7 s^−1^. TPK3 was initially thought to be a voltage-dependent potassium channel fulfilling this role [27]. However, it was found to localize to the tonoplast [28]; therefore, within the model structure, another transport pathway is required to move potassium from the lumen to the stroma and balance potassium import by KEA3. The 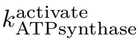 optimum lies at 0.001 s^−^1, being 10 fold lower as in [74]. This could be explained by the additional data we used, such as gas assimilation, which constrained the model differently than using only NPQ data. *pK*_reg_, the parameter representing the *pK*_a_ of the b6f side chain responsible for photosynthetic control, was also well identifiable by the data to 6.3. The profile shows that shifting it slightly higher to 6.44 produces a Λ_*i*_(p) of 742, which represents a 50 % increase compared to the optimum, suggesting that a *pK*_a_ around 6.3 is essential to reproduce the data.

The sensitivity analysis places these parameters in context. The PMF was most sensitive to the total chloride ion concentration and to photosynthetic control in high light. Total chloride ion concentration can be so impactful because our fit determined it to be 30 mM, which can be the limiting factor if it varies by a factor of two. Increasing ion concentration decreases overall PMF in high light because more chloride ions transferred into the lumen would reduce the membrane potential. The model shows that reduced electrostatic forces on protons are evident in low light, where the delta pH is elevated. Chloride ions might be crucial because their total concentration limits ion transport. For potassium, the opposite might be true, as the fit indicated a higher total concentration. Here, membrane ion transport might be limiting, as the Sobol’ analysis suggests that varying the opening thresholds of KEA3 and the unknown channel strongly affects PMF.

Photosynthetic control, represented by the parameter *pK*_reg_, showed the strongest impact on PMF in high light. With an increase in *pK*_a_ to 6.47, photosynthetic control would begin at higher pH_lumen_, decreasing PMF by about 25 mV. With a *pK*_a_ of 6.8, total PMF is decreased by about 110 mV. This aligns with what Jahns *et al*. found [75], where they used a mutant of the Rieske subunit which shifted the *pK*_a_ of photosynthetic control up by one pH unit to investigate photosynthetic control. The mutant did not allow a larger ΔpH than two to form. In our predicted PMF partitioning, with a *pK*_a_ of 6.8, the ΔpH is downregulated to 110 mV, corresponding to 1.69 pH units.

While the amounts of PSII and PSI and the catalytic capacities of CBB cycle enzymes strongly affected PMF under low-light conditions, consistent with PMF linking the supply of ATP and NADPH to their demand [2, 3], ion transport mainly affected PMF during light transitions. The high sensitivity of the PMF to the amounts of PSII and PSI is notable because, in the model, these amounts are constant, whereas in the leaf the number of active and connected photosystems changes on the timescales of our protocols. Light-induced reorganization of grana stacks alters the connectivity between PSII, PSI and the plastoquinone pool [76, 77], photoinhibition reduces the number of active PSII, and changes in lumen volume alter proton buffering and the relationship between translocated charge and ΔΨ [78]. Our results therefore suggest that such structural dynamics of the thylakoid could be an important, so far unmodeled, determinant of PMF size and composition.

The model also predicts how the PMF is partitioned, the quantity at the center of the debate between ECS-based estimates of a substantial ΔΨ [19, 12] and studies arguing for a ΔpH-dominated PMF [29, 30]. Our simulations support a light-dependent partitioning: ΔpH dominated the PMF in high light, whereas ΔΨ contributed about half in low light (Fig. 7). Consistent with Davis *et al*.[21], ion transport shaped the partitioning mainly at light transitions, when ΔΨ and ΔpH changed transiently. Because the ECS was fitted after affine scaling, the predicted partitioning depends on the model’s fixed charge-to-voltage conversion and lumen buffer capacity, making independent quantification of these parameters the most direct test of this.

Our conclusions are naturally subject to limitations of the model structure, the data and the inference methods, which we discuss in turn. Like any model, ours simplifies several processes. Photorespiration is not included, although at 400 ppm CO_2_ it contributes substantially to the ATP demand [76], so fitted parameters of the CBB cycle, the ATP synthase and CEF may partly compensate for its absence. The intercellular CO_2_ concentration was kept constant, so stomatal and mesophyll conductance dynamics are not captured, although C_*i*_ can be varied dynamically in simulations. Redox activation of the ATP synthase is approximated by a light-dependent switch, and NPQ by a phenomenological four-state quencher. Two of the five ion transport pathways are hypothetical: their well-identified rate constants indicate that the model requires ion fluxes of this magnitude, not that dedicated channels exist, and transporters with overlapping roles, such as ClCe and the hypothetical Cl^−^ channel, cannot be fully distinguished. Furthermore, we deliberately used data from a single study [51] to avoid inconsistencies between measurements, which restricts our analysis to one genotype, one growth condition, and one CO_2_ concentration. For the ECS, we used a single replicate and mapped the simulated ΔΨ onto the data by an affine transformation, and deriving ΔΨ from the ECS assumes that the absorbance change is exclusively electrochromic, an assumption that has been questioned [29, 30].

Hence, the natural next step is to test the model against independent data, such as measurements under other light regimes and CO_2_ concentrations, or on mutants lacking specific ion transporters, and then to use it to systematically examine how the PMF is partitioned between ΔΨ and ΔpH across conditions. Such tests need not be restricted to our own data: the model is available in the Greensloth database [61], where users can import their own measurements and fit the model directly in the browser, so that it can be evaluated independently against data from other laboratories, species, and growth conditions. Our analysis further suggests which type of measurement would be most informative. The three measurements used here were recorded on separate samples and constrain the model only jointly, and the predicted PMF partitioning depends on parameters, such as the lumen buffer capacity, that none of them resolve. Instruments that record two observables simultaneously in the same sample, such as chlorophyll fluorescence together with the signal of genetically encoded pH sensors [79], would remove variation between samples and allow predicted relations between variables to be tested within a single measurement. For example, the relation between the redox state of the plastoquinone pool and lumen pH underlies photosynthetic control, the process that dominated the PMF in high light in our analysis. Coupling kinetic models with structural information, for example from cryo-electron tomography [80, 77], may further be required to capture the role of thylakoid structure. Existing measurements can thus reveal the mechanisms that shape PMF dynamics, provided they are combined, but inferring PMF magnitude also requires knowledge of the photosynthetic machinery that sets its capacity.

## Author contributions

JAE: Conceptualization, Methodology, Software, Formal analysis, Investigation, Visualization, Writing – original draft, Writing – review & editing. AM: Conceptualization, Methodology, Supervision, Funding acquisition, Project administration, Writing – original draft, Writing – review & editing.

## Acknowledgment

We thank Thekla von Bismarck and Ute Armbruster for providing the experimental data necessary for this project. We are grateful to Quang Huy Nguyen for insightful discussions on proton motive force, to Mar-vin van Aalst for his work on accelerating the simulations, and to Tim Nies for careful proofreading of the manuscript.

## References

1. Matthew P. Johnson. Photosynthesis. Essays in Biochemistry, 60(3):255–273, 10 2016.

2. David M. Kramer and John R. Evans. The importance of energy balance in improving photosynthetic productivity. Plant Physiology, 155(1):70–78, 01 2011.

3. Anna Matuszyńska, Nima P Saadat, and Oliver Ebenhöh. Balancing energy supply during photosynthesis - a theoretical perspective. Physiol. Plant., 166(1):392–402, May 2019.

4. Zsofia R. Stangl, Lasse Tarvainen, Göran Wallin, and John D. Marshall. Limits to photosynthesis: seasonal shifts in supply and demand for co2 in scots pine. New Phytologist, 233(3):1108–1120, 2022.

5. Sabeeha Merchant and Michael R. Sawaya. The light reactions: A guide to recent acquisitions for the picture gallery. The Plant Cell, 17(3):648–663, 03 2005.

6. Alexandrina Stirbet, Sushma Naithani, Dmitry Shevela, Lars Björn, Julian Eaton-Rye, and Arthur Nonomura. On the reactions in oxygenic photosynthesis as related to the z-scheme: Contributions by govindjee. LS-International Journal of Life Sciences, 12:151–185, 01 2023.

7. Klaus Brettel and Winfried Leibl. Electron transfer in photosystem i. Biochimica et Biophysica Acta (BBA) - Bioenergetics, 1507(1):100–114, 2001.

8. Tanai Cardona, Arezki Sedoud, Nicholas Cox, and A. William Rutherford. Charge separation in photosystem ii: A comparative and evolutionary overview. Biochimica et Biophysica Acta (BBA) - Bioenergetics, 1817(1):26–43, 2012.

9. Mahir Mamedov, Govindjee, Victor Nadtochenko, and Alexey Semenov. Primary electron transfer processes in photosynthetic reaction centers from oxygenic organisms. Photosynthesis Research, 125(1):51–63, Aug 2015.

10. Alexander Hahn, Janet Vonck, Deryck J. Mills, Thomas Meier, and Werner Kühlbrandt. Structure, mechanism, and regulation of the chloroplast atp synthase. Science, 360(6389):eaat4318, 2018.

11. Atsuko Kanazawa, Peter Neofotis, Geoffry A. Davis, Nicholas Fisher, and David M. Kramer. Diversity in photoprotection and energy balancing in terrestrial and aquatic phototrophs. In Anthony W.D. Larkum, Arthur R. Grossman, and John A. Raven, editors, Photosynthesis in Algae: Biochemical and Physiological Mechanisms, pages 299–327. Springer International Publishing, Cham, 2020.

12. David M. Kramer, Jeffrey A. Cruz, and Atsuko Kanazawa. Balancing the central roles of the thylakoid proton gradient. Trends in Plant Science, 8(1):27–32, Jan 2003.

13. Graham Noctor and Christine H. Foyer. A re-evaluation of the atp: nadph budget during c3 photosynthesis: a contribution from nitrate assimilation and its associated respiratory activity? Journal of Experimental Botany, 49(329):1895–1908, 12 1998.

14. Yuri Munekage, Mihoko Hashimoto, Chikahiro Miyake, Ken-Ichi Tomizawa, Tsuyoshi Endo, Masao Tasaka, and Toshiharu Shikanai. Cyclic electron flow around photosystem i is essential for photosynthesis. Nature, 429(6991):579–582, Jun 2004.

15. Kozi Asada. The water-water cycle in chloroplasts: scavenging of active oxygens and dissipation of excess photons. Annual Review of Plant Physiology and Plant Molecular Biology, 50:601–639, 1999.

16. Marjaana Suorsa. Cyclic electron flow provides acclimatory plasticity for the photosynthetic machinery under various environmental conditions and developmental stages. Frontiers in Plant Science, Volume 6 – 2015, 2015.

17. Deserah D. Strand, Nicholas Fisher, and David M. Kramer. The higher plant plastid nad(p)h dehydrogenase-like complex (ndh) is a high efficiency proton pump that increases atp production by cyclic electron flow. Journal of Biological Chemistry, 292(28):11850–11860, 2017.

18. Geoffry A Davis, Atsuko Kanazawa, Mark Aurel Schöttler, Kaori Kohzuma, John E Froehlich, A. William Rutherford, Mio Satoh-Cruz, Deepika Minhas, Stefanie Tietz, Amit Dhingra, and David M Kramer. Limitations to photosynthesis by proton motive force-induced photosystem ii photodamage. eLife, 5:e16921, oct 2016.

19. Jeffrey A. Cruz, Colette A. Sacksteder, Atsuko Kanazawa, and David M. Kramer. Contribution of electric field (Δψ) to steady-state transthylakoid proton motive force (pmf) in vitro and in vivo. control of pmf parsing into Δψ and Δph by ionic strength. Biochemistry, 40(5):1226–1237, 02 2001.

20. Hans-Henning Kunz, Ute Armbruster, Susanne Mühlbauer, Jan de Vries, and Geoffry A. Davis. Chloroplast ion homeostasis – what do we know and where should we go? New Phytologist, 243(2):543–559, 2024.

21. Geoffry A Davis, A. William Rutherford, and David M Kramer. Hacking the thylakoid proton motive force for improved photosynthesis: modulating ion flux rates that control proton motive force partitioning into Δψ and ΔpH. Philos. Trans. R. Soc. Lond. B Biol. Sci., 372(1730):20160381, September 2017.

22. Andrei Herdean, Enrico Teardo, Anders K Nilsson, Bernard E Pfeil, Oskar N Johansson, Renáta Ünnep, Gergely Nagy, Ottó Zsiros Somnath Dana, Katalin Solymosi, Győző Garab Ildikó Szabó, Cornelia Spetea, and Björn Lundin. A voltage-dependent chloride channel fine-tunes photosynthesis in plants. Nat. Commun., 7(1):11654, May 2016.

23. Andrei Herdean, Hugues Nziengui, Ottó Zsiros Katalin Solymosi, Győző Garab Björn Lundin, and Cornelia Spetea. The arabidopsis thylakoid chloride channel AtCLCe functions in chloride homeostasis and regulation of photosynthetic electron transport. Front. Plant Sci., 7:115, February 2016.

24. Emilija Dukic, Peter J Gollan, Steffen Grebe, Virpi Paakkarinen, Andrei Herdean, Eva-Mari Aro, and Cornelia Spetea. The arabidopsis thylakoid chloride channel ClCe regulates ATP availability for light-harvesting complex II protein phosphorylation. Front. Plant Sci., 13:1050355, November 2022.

25. Ute Armbruster, L Ruby Carrillo, Kees Venema, Lazar Pavlovic, Elisabeth Schmidtmann, Ari Kornfeld, Peter Jahns, Joseph A Berry, David M Kramer, and Martin C Jonikas. Ion antiport accelerates photosynthetic acclimation in fluctuating light environments. Nat. Commun., 5(1):5439, November 2014.

26. Michał Uflewski Tobias Rindfleisch, Kübra Korkmaz, Enrico Tietz, Sarah Mielke, Viviana Correa Galvis, Beatrix Dünschede, Marcin Luzarowski, Aleksandra Skirycz, Markus Schwarzländer, Deserah D. Strand, Alexander P. Hertle, Danja Schünemann, Dirk Walther, Anja Thalhammer, Martin Wolff, and Ute Armbruster. The thylakoid proton antiporter kea3 regulates photosynthesis in response to the chloroplast energy status. Nature Communications, 15(1):2792, Mar 2024.

27. Luca Carraretto, Elide Formentin, Enrico Teardo, Vanessa Checchetto, Martino Tomizioli, Tomas Morosinotto, Giorgio Mario Giacometti, Giovanni Finazzi, and Ildikó Szabó. A thylakoid-located two-pore k<sup>+</sup> channel controls photosynthetic light utilization in plants. Science, 342(6154):114–118, 2013.

28. Ricarda Höhner, Viviana Correa Galvis, Deserah D. Strand, Carsten Völkner, Moritz Krämer, Michaela Messer, Firdevs Dinc, Inga Sjuts, Bettina Bölter, David M. Kramer, Ute Armbruster, and Hans-Henning Kunz. Photosynthesis in arabidopsis is unaffected by the function of the vacuolar k+ channel tpk3. Plant Physiology, 180(3):1322–1335, 07 2019.

29. Matthew P. Johnson and Alexander V. Ruban. Rethinking the existence of a steady-state δψ component of the proton motive force across plant thylakoid membranes. Photosynthesis Research, 119(1):233–242, Feb 2014.

30. Sam Wilson, Matthew P. Johnson, and Alexander V. Ruban. Proton motive force in plant photosynthesis dominated by Δph in both low and high light. Plant Physiology, 187(1):263–275, 09 2021.

31. Erik H. Murchie and Krishna K. Niyogi. Manipulation of photoprotection to improve plant photosynthesis. Plant Physiology, 155(1):86–92, 01 2011.

32. Ginga Shimakawa and Chikahiro Miyake. Oxidation of p700 ensures robust photosynthesis. Frontiers in Plant Science, Volume 9 – 2018, 2018.

33. Barbara Demmig-Adams and William W. Adams. Photoprotection and other responses of plants to high light stress. Annual Review of Plant Biology, 43(Volume 43, 1992):599–626, 1992.

34. Alexander V. Ruban and Matthew P. Johnson. Xanthophylls as modulators of membrane protein function. Archives of Biochemistry and Biophysics, 504(1):78–85, 2010.

35. Gero Miesenböck, Dino A. De Angelis, and James E. Rothman. Visualizing secretion and synaptic transmission with ph-sensitive green fluorescent proteins. Nature, 394(6689):192–195, Jul 1998.

36. Hong Yang, Xiaojun Pu, Lu Wang, Li Liu, and Steven M. Theg. A new fluorescence-based method to monitor the ph in the thylakoid lumen using gfp variants. Biochemical and Biophysical Research Communications, 486(1):1–5, 2017.

37. Jan-Ole Niemeier, Pedro Barreto, Bruce Morgan, and Markus Schwarzländer. In vivo monitoring of energy metabolism with genetically encoded fluorescent biosensors. Annual Review of Plant Biology, 77(Volume 77, 2026):641–675, 2026.

38. Neil R. Baker. Chlorophyll Fluorescence: A Probe of Photosynthesis In Vivo. Annual Review of Plant Biology, 59(1):89–113, June 2008.

39. Horst Tobias Witt. Energy conversion in the functional membrane of photosynthesis. analysis by light pulse and electric pulse methods: The central role of the electric field. Biochimica et Biophysica Acta (BBA) - Reviews on Bioenergetics, 505(3):355–427, 1979.

40. Benjamin Bailleul, Pierre Cardol, Cecile Breyton, and Giovanni Finazzi. Electrochromism: a useful probe to study algal photosynthesis. Photosynthesis Research, 106(1-2), 2010.

41. Kenji Takizawa, Jeffrey A. Cruz, Atsuko Kanazawa, and David M. Kramer. The thylakoid proton motive force in vivo. quantitative, non-invasive probes, energetics, and regulatory consequences of light-induced pmf. Biochimica et Biophysica Acta (BBA) - Bioenergetics, 1767(10):1233–1244, 2007.

42. Colette A. Sacksteder, Atsuko Kanazawa, Michael E. Jacoby, and David M. Kramer. The proton to electron stoichiometry of steady-state photosynthesis in living plants: A proton-pumping q cycle is continuously engaged. Proceedings of the National Academy of Sciences, 97(26):14283–14288, 2000.

43. Thomas D. Sharkey and Yuan Xu. arbon dioxide release during photosynthesis: Connecting gas exchange behavior with biochemistry. Plant, Cell & Environment, 49(1):309–319, 2026.

44. Bernard Genty, Jeremy Harbinson, Jean-Marie Briantais, and Neil R. Baker. The relationship between non-photochemical quenching of chlorophyll fluorescence and the rate of photosystem 2 photochemistry in leaves. Photosynthesis Research, 25(3):249–257, Sep 1990.

45. Andreas Raue, Clemens Kreutz, Thomas Maiwald, Julie Bachmann, Marcel Schilling, Ursula Klingmüller, and Jens Timmer. Structural and practical identifiability analysis of partially observed dynamical models by exploiting the profile likelihood. Bioinformatics, 25(15):1923–1929, 08 2009.

46. Jennifer E. Johnson and Joseph A. Berry. The role of cytochrome b6f in the control of steady-state photosynthesis: a conceptual and quantitative model. Photosynthesis Research, 148(3):101–136, Jun 2021.

47. Meng Li, Vaclav Svoboda, Geoffry Davis, David Kramer, Hans-Henning Kunz, and Helmut Kirchhoff. Impact of ion fluxes across thylakoid membranes on photosynthetic electron transport and photoprotection. Nat. Plants, 7(7):979–988, July 2021.

48. Alejandro Morales, Xinyou Yin, Jeremy Harbinson, Steven M. Driever, Jaap Molenaar, David M. Kramer, and Paul C. Struik. In silico analysis of the regulation of the photosynthetic electron transport chain in c3 plants. Plant Physiology, 176(2):1247–1261, 02 2018.

49. Nima P. Saadat, Tim Nies, Marvin van Aalst, Brandon Hank, Büsra Demirtas, Oliver Ebenhöh, and Anna Matuszyńska. Computational analysis of alternative photosynthetic electron flows linked with oxidative stress. Front. Plant Sci., 12:750580, 2021.

50. Lianhong Gu, Stephen G. Pallardy, Kevin Tu, Beverly E. Law, and Stan D. Wullschleger. Reliable estimation of biochemical parameters from c3 leaf photosynthesis–intercellular carbon dioxide response curves. Plant, Cell & Environment, 33(11):1852–1874, 2010.

51. Thekla von Bismarck, Kübra Korkmaz, Jeremy Ruß, Kira Skurk, Elias Kaiser, Viviana Correa Galvis, Jeffrey A. Cruz, Deserah D. Strand, Karin Köhl, Jürgen Eirich, Iris Finkemeier, Peter Jahns, David M. Kramer, and Ute Armbruster. Light acclimation interacts with thylakoid ion transport to govern the dynamics of photosynthesis in arabidopsis. New Phytologist, 237(1):160–176, 2023.

52. David M. Kramer, Colette A. Sacksteder, and Jeffrey A. Cruz. How acidic is the lumen? Photosynthesis Research, 60:151–163, 1999.

53. Andre Potthoff. The perks of pirks and dirks mechanistic modelling of photosynthetic relaxation kinetics under rapid light protocols. Bachelor’s thesis, Heinrich Heine University Düsseldorf, 2025. Unpublished.

54. Oliver Ebenhöh, Geoffrey Fucile, Giovanni Finazzi, Jean-David Rochaix, and Michel Goldschmidt-Clermont. Short-term acclimation of the photosynthetic electron transfer chain to changing light: a mathematical model. Philosophical Transactions of the Royal Society B: Biological Sciences, 369(1640):20130223, 04 2014.

55. Karl Werdan, Hans W. Heldt, and Mirjana Milovancev. The role of ph in the regulation of carbon fixation in the chloroplast stroma. studies on co2 fixation in the light and dark. Biochimica et Biophysica Acta (BBA) - Bioenergetics, 396(2):276–292, 1975.

56. Ulrike Junesch and Peter Gräber. Influence of the redox state and the activation of the chloroplast atp synthase on proton-transport-coupled atp synthesis/hydrolysis. Biochimica et Biophysica Acta (BBA) - Bioenergetics, 893(2):275–288, 1987.

57. Keisuke Yoshida, Ayaka Hara, Kazunori Sugiura, Yuki Fukaya, and Toru Hisabori. Thioredoxin-like2/2-cys peroxiredoxin redox cascade supports oxidative thiol modulation in chloroplasts. Proceedings of the National Academy of Sciences, 115(35):E8296–E8304, 2018.

58. Marvin van Aalst, Tim Nies, Tobias Pfennig, and Anna Matuszyńska. MxlPy—python package for mechanistic learning and hybrid modelling in life science. Bioinformatics Advances, 5(1):vbaf294, 2026.

59. Ernst Hairer and Gerhard Wanner. Stiff differential equations solved by radau methods. Journal of Computational and Applied Mathematics, 111(1):93–111, 1999.

60. Marvin van Aalst, Alienor Lahlou, Tanvir Hassan, William Gaultier, David Colliaux, and Anna Matuszyńska. Web-based collaborative model development in interdisciplinary consortia: Design principles and practical guidance. PLoS Biol., 24(6):e3003825, June 2026.

61. Elouen Corvest, Marvin van Aalst, Tim Nies, Quang Huy Nguyen, Josha Ebeling, Maja Strauch, El Hadji Malick Cisse, Tanvir Hassan, and Anna Matuszyńska. Greensloth: a curated database and executable platform for mechanistic photosynthesis models. bioRxiv, 2026.

62. Pauli Virtanen, Ralf Gommers, Travis E. Oliphant, Matt Haberland, Tyler Reddy, David Cournapeau, Evgeni Burovski, Pearu Peterson, Warren Weckesser, Jonathan Bright, Stéfan J. van der Walt, Matthew Brett, Joshua Wilson, K. Jarrod Millman, Nikolay Mayorov, Andrew R. J. Nelson, Eric Jones, Robert Kern, Eric Larson,J J Carey, İIlhan Polat, Yu Feng, Eric W. Moore, Jake VanderPlas, Denis Laxalde, Josef Perktold, Robert Cimrman, Ian Henriksen, E. A. Quintero, Charles R. Harris, Anne M. Archibald, Antônio H. Ribeiro, Fabian Pedregosa, Paul van Mulbregt, and SciPy 1.0 Contributors. SciPy 1.0: Fundamental Algorithms for Scientific Computing in Python. Nature Methods, 17:261–272, 2020.

63. Yannik Schälte, Fabian Fröhlich, Paul J Jost, Jakob Vanhoefer, Dilan Pathirana, Paul Stapor, Polina Lakrisenko, Dantong Wang, Elba Raimúndez, Simon Merkt, Leonard Schmiester, Philipp Städter, Stephan Grein, Erika Dudkin, Domagoj Doresic, Daniel Weindl, and Jan Hasenauer. pypesto: a modular and scalable tool for parameter estimation for dynamic models. Bioinformatics, 39(11):btad711. 11 2023.

64. Jon Herman and Will Usher. SALib: An open-source python library for sensitivity analysis. The Journal of Open Source Software, 2(9), jan 2017.

65. Christopher C. Hall, Jeffrey Cruz, Magnus Wood, Robert Zegarac, Dustin DeMars, Joel Carpenter, Atsuko Kanazawa, and David Kramer. Photosynthetic measurements with the idea spec: an integrated diode emitter array spectrophotometer/fluorometer. In Photosynthesis Research for Food, Fuel and the Future, pages 184–188, Berlin, Heidelberg, 2013. Springer Berlin Heidelberg.

66. Ulrich Schreiber. Pulse-amplitude-modulation (pam) fluorometry and saturation pulse method: An overview. In George Christos Papageorgiou and Govindjee, editors, Chlorophyll a Fluorescence: A Signature of Photosynthesis, pages 279–319. Springer Netherlands, Dordrecht, 2004.

67. Wolfgang Bilger and Olle Björkman. Role of the xanthophyll cycle in photoprotection elucidated by measurements of light-induced absorbance changes, fluorescence and photosynthesis in leaves of hedera canariensis. Photosynth. Res., 25(3):173–185, September 1990.

68. Edward B Lochocki and Justin M McGrath. Widely used variants of the farquhar-von-caemmerer-berry model can cause errors in parameter estimation. in silico Plants, 7(2):diaf014. 07 2025.

69. Graham D. Farquhar, Susanne von Caemmerer, and Joseph A. Berry. A biochemical model of photosynthetic co2 assimilation in leaves of c3 species. Planta, 149(1):78–90, Jun 1980.

70. Charlie Mathiot and Jean Alric. Standard units for electrochromic shift measurements in plant biology. Journal of Experimental Botany, 72(18):6467–6473, 09 2021.

71. Clemens Kreutz, Andreas Raue, Daniel Kaschek, and Jens Timmer. Profile likelihood in systems biology. The FEBS Journal, 280(11):2564–2571, 2013.

72. Andrea Saltelli, Paola Annoni, Ivano Azzini, Francesca Campolongo, Marco Ratto, and Stefano Tarantola. Variance based sensitivity analysis of model output. design and estimator for the total sensitivity index. Computer Physics Communications, 181(2):259–270, 2010.

73. Ilya M. Sobol. Global sensitivity indices for nonlinear mathematical models and their monte carlo estimates. Mathematics and Computers in Simulation, 55(1):271–280, 2001.

74. Anna Matuszyńska, Somayyeh Heidari, Peter Jahns, and Oliver Ebenhöh. A mathematical model of non-photochemical quenching to study short-term light memory in plants. Biochimica et Biophysica Acta (BBA) - Bioenergetics, 1857(12):1860–1869, 2016.

75. Peter Jahns, Maria Graf, Yuri Munekage, and Toshiharu Shikanai. Single point mutation in the rieske iron–sulfur subunit of cytochrome b 6/f leads to an altered ph dependence of plastoquinol oxidation in arabidopsis. FEBS Letters, 519:99–102, 04 2002.

76. Berkley J. Walker, Andy VanLoocke, Carl J. Bernacchi, and Donald R. Ort. The costs of photorespiration to food production now and in the future. Annual Review of Plant Biology, 67(Volume 67, 2016):107–129, 2016.

77. Wojciech Wietrzynski, Lorenz Lamm, William HJ Wood, Matina-Jasemi Loukeri, Lorna Malone, Tingying Peng, Matthew P Johnson, and Benjamin D Engel. Molecular architecture of thylakoid membranes within intact spinach chloroplasts. eLife, 14:RP105496, sep 2025.

78. Helmut Kirchhoff, Chris Hall, Magnus Wood, Miroslava Herbstová, Onie Tsabari, Reinat Nevo, Dana Charuvi, Eyal Shimoni, and Ziv Reich. Dynamic control of protein diffusion within the granal thylakoid lumen. Proceedings of the National Academy of Sciences, 108(50):20248–20253, 2011.

79. Sanchali Nanda, Tatyana Shutova, Maximiliano Cainzos, Chen Hu, Bart Sasbrink, Pushan Bag, Tristan den Blanken, Ronald Buijs, Lex van der Gracht, Frans Hendriks, Petar Lambrev, Rob Limburg, Vincenzo Mascoli, Wojciech J Nawrocki, Michael Reus, Ramon Parmessar, Björn Singerling, Ivo H M Stokkum, Stefan Jansson, and Alfred R. Holzwarth. Chlorospec: A new in vivo chlorophyll fluorescence spectrometer for simultaneous wavelength- and time-resolved detection. Physiologia Plantarum, 176(2):e14306, 2024.

80. Benjamin D Engel, Miroslava Schaffer, Luis Kuhn Cuellar, Elizabeth Villa, Jürgen M Plitzko, and Wolfgang Baumeister. Native architecture of the Chlamydomonas chloroplast revealed by in situ cryo-electron tomography. eLife, 4:e04889. jan 2015.

