## Supplementary material for "Can we infer mechanisms governing PMF generation and partitioning from existing experimental data? A model-based identifiability analysis": Sup

November 23, 2026

**Supplementary Material for the manuscript by Ebeling and Matuszyńska titled "Can we infer mechanisms governing PMF generation and partitioning from existing experimental data? A model-based identifiability analysis."**

#### 1 Supplementary Figures

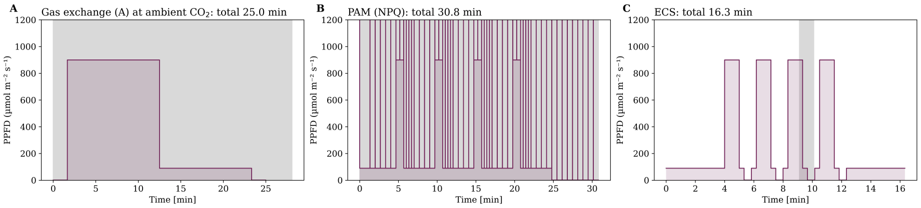


Figure S1: Measurement light protocol of gas exchange, SP-PAM, and ECS. Magenta shaded areas highlight the PPFD during the protocol, while the shaded area denotes the timeframe used for fitting.

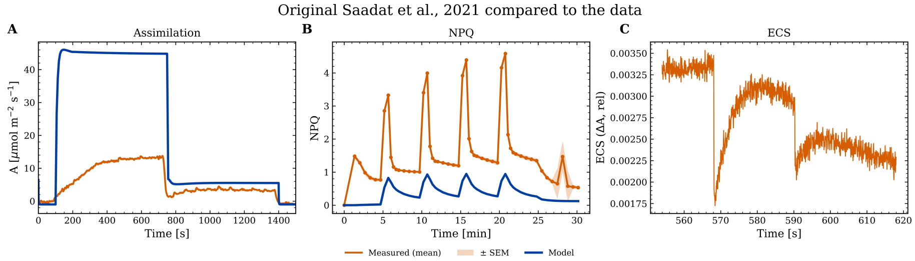


Figure S2: Comparison of the Saadat et al. (2021) model with the measurement data used in the study, highlighting the improvements made.

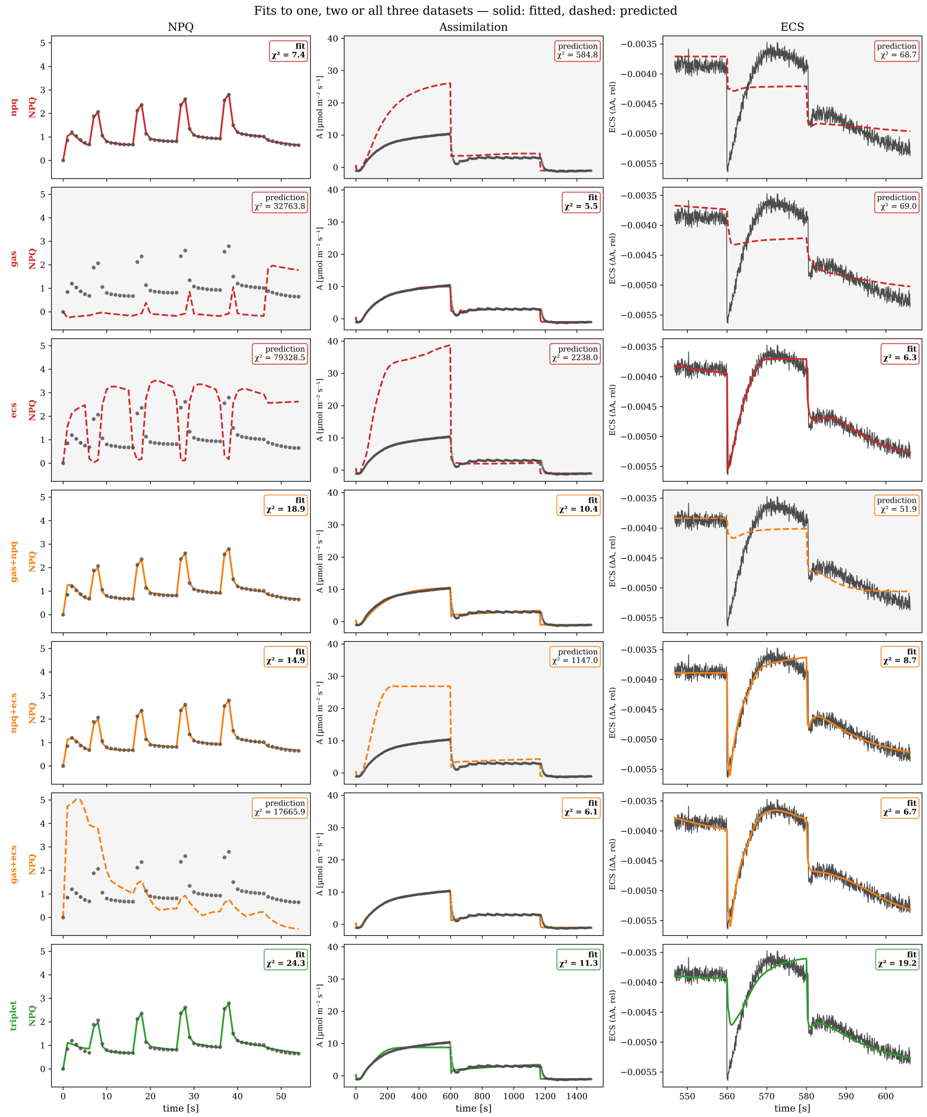


Figure S3: Results of the progressive fitting, plotting the simulations (fits as solid lines and predictions as dashed) against the experimental data for each phase of the progressive fitting.

### Model information

#### 1.1 Modified, added or removed parameters

Table S1: Model parameters that differ between Saadat et al. (2021) and this study. Fitted values are given at the final optimum.

| Parameter | Saadat 2021 | This study | Fitting bounds | Parameter category |
|---|---|---|---|---|
| $[CO_2]$ | 0.20 | 0.0136 | – | Literature-derived; atmospheric $CO_2$ condition |
| $b_{\mathrm{H}}$ | 100.00 | 200.00 | – | Assumed |
| $b_{\mathrm{H}}^{\mathrm{stroma}}$ | – | 500.00 | – | Literature-derived / assumed; larger buffering than in the lumen, chosen so that stromal pH stays between 7 and 8 |
| $V_{\mathrm{charge}}$ | – | 0.00077 | – | Assumed; keeps the membrane potential in the reported physiological range |
| $\mathrm{pH}_{\mathrm{stroma}}$ | 7.90 | – | – | Removed with dynamic stromal pH |
| $[\mathrm{H}^+_{\mathrm{stroma}}]$ | 0.000013 | – | – | Removed with dynamic stromal pH |
| $k_{\mathrm{VCCN1}}$ | – | 3.01 | 0.10–100.00 | Fitted |
| $\mathrm{VCCN1}_{\mathrm{threshold}}^{\Delta\psi}$ | – | 0.0044 | 0.0010–0.10 | Fitted |
| $k_{\mathrm{ClCe}}$ | – | 1.00 | 0.01–1.00 | Fitted |
| $\mathrm{ClCe}_{\mathrm{threshold}}^{\mathrm{ATP}}$ | – | 0.0358 | 0.0010–0.50 | Fitted |
| $k_{\mathrm{Cl\,leak}}$ | – | 3.28 | 0.10–100.00 | Fitted |
| $\mathrm{PQ}_{\mathrm{thr}}^{\mathrm{Cl\,leak}}$ | – | 6.61 | 6.00–15.00 | Fitted; bounds based on assumed activity of an unknown channel |
| $perm_{\mathrm{K+}}$ | – | 5.69 | 0.10–100.00 | Fitted |
| $\mathrm{K}_{\mathrm{threshold}}^{+\Delta\psi}$ | – | 0.0526 | 0.0010–0.10 | Fitted |
| $k_{\mathrm{KEA3}}$ | – | 17.86 | 10.00–100.00 | Fitted |
| $\mathrm{KEA3}_{\mathrm{pHreg}}^{\mathrm{KEA3}}$ | – | 7.62 | 7.40–8.00 | Fitted; bounds based on reported activity of KEA3 |
| $\mathrm{KEA3}_{\mathrm{threshold}}^{\mathrm{ATP}}$ | – | 0.01 | – | Assumed / literature-derived, based on reported activity of KEA3 |
| $[\mathrm{K}^+]_{\mathrm{tot}}$ | – | 68.97 | 25.00–100.00 | Fitted |
| $[\mathrm{Cl}^-]_{\mathrm{tot}}$ | – | 15.65 | 1.00–50.00 | Fitted |
| $k_{\mathrm{ATPsynthase}}$ | – | 20.00 | – | Assumed |
| $k_{\mathrm{ATPsynthase}}^{\mathrm{activate}}$ | – | 0.0012 | 0.00010–1.00 | Fitted |
| $k_{\mathrm{ATPsynthase}}^{\mathrm{deactivate}}$ | – | 0.00010 | 0.00010–0.10 | Fitted |
| $\mathrm{b_6f_{tot}}$ | – | 1.00 | – | Literature-derived |

*Continued on next page*

Table S1: Model parameters (continued)

| Parameter | Saadat 2021 | This study | Fitting bounds | Parameter category |
|---|---|---|---|---|
| $v_{\mathrm{b_6f}}^{\mathrm{max}}$ | – | 300.00 | – | Literature-derived / assumed |
| $n_{\mathrm{H}}^{\mathrm{b_6f}}$ | – | 3.00 | – | Assumed |
| $\mathrm{p}K_{\mathrm{reg}}$ | – | 6.30 | 6.30–7.00 | Fitted |
| $k_{\mathrm{H}}^{\mathrm{Qslope}}$ | – | $5.00 \times 10^{9}$ | – | Added with dynamic photosystems / literature-derived |
| $k_{\mathrm{NDH}}$ | – | 6.97 | 0.01–10.00 | Fitted |
| $k_{\mathrm{b_6f}}^{\mathrm{cat}}$ | 2.50 | – | – | Removed with the $\mathrm{b_6f}$ change |
| $k_{\mathrm{FNR,PSI}}^{\mathrm{cat}}$ | $2.50 \times 10^{5}$ | – | – | Removed with dynamic photosystems |
| $E_{0,\mathrm{FNR,PSI}}$ | 1.00 | – | – | Removed with dynamic photosystems |
| $\gamma_0$ | 0.10 | 0.0677 | 0.0010–1.00 | Fitted |
| $\gamma_1$ | 0.25 | 0.94 | 0.0010–10.00 | Fitted |
| $\gamma_2$ | 0.60 | 3.94 | 0.0010–10.00 | Fitted |
| $\gamma_3$ | 0.15 | 3.41 | 0.0010–10.00 | Fitted |
| $k_{\mathrm{LHC}}^{\mathrm{prot}}$ | 0.0096 | 0.33 | 0.0010–1.00 | Fitted |
| $k_{\mathrm{LHC}}^{\mathrm{deprot}}$ | 0.0096 | 0.0756 | 0.0010–1.00 | Fitted |
| $n_{\mathrm{H}}^{\mathrm{LHC}}$ | 3.00 | 2.16 | 1.00–20.00 | Fitted |
| $k_{\mathrm{LHC\ protonation}}^{\mathrm{sat}}$ | 5.80 | 5.50 | 5.50–6.50 | Fitted |
| $k_{\mathrm{VDE}}$ | 0.0024 | 0.00013 | 0.000010–1.00 | Fitted |
| $n_{\mathrm{H}}^{\mathrm{VDE}}$ | 5.00 | 19.89 | 1.00–20.00 | Fitted |
| $k_{\mathrm{VDE}}^{\mathrm{sat}}$ | 5.80 | 6.40 | 5.50–6.50 | Fitted |
| $k_{\mathrm{ZE}}$ | 0.00024 | 0.00060 | 0.000010–1.00 | Fitted |
| $k_{\mathrm{FTR}}$ | 1.00 | 0.13 | 0.0010–5.00 | Fitted |
| $k_{\mathrm{TrxActivation}}^{\mathrm{f}}$ | 1.00 | 0.0037 | 0.000010–0.10 | Fitted |
| $k_{\mathrm{TrxInactivation}}^{\mathrm{f}}$ | 0.10 | 0.0118 | 0.000010–1.00 | Fitted |
| $k2_{\mathrm{inactivation}}^{\mathrm{Trx}}$ | – | 0.0238 | 0.0010–1.00 | Fitted |

#### 1.2 Ordinary differential equations

Table S2: Ordinary differential equations of the model. $v$ denotes reaction rates.

| Variable | Differential equation |
|---|---|
| $d[3PGA]/dt$ | $= -v^{ex}_{3PGA} + 2\,v_{Rubisco} - \mathbf{x_0}$ |
| $d[BPGA]/dt$ | $= -v_{GAPDH} + \mathbf{x_0}$ |
| $d[GAP]/dt$ | $= -v_{ALD,FBP} - v^{ex}_{GAP} + v_{GAPDH} - \mathbf{x_0} - \mathbf{x_1} - \mathbf{x_2}$ |
| $d[DHAP]/dt$ | $= -v_{ALD,SBP} - v_{ALD,FBP} - v^{ex}_{DHAP} + \mathbf{x_0}$ |
| $d[FBP]/dt$ | $= v_{ALD,FBP} - v_{FBPase}$ |
| $d[F6P]/dt$ | $= v_{FBPase} - v_{G6PI} - \mathbf{x_0}$ |
| $d[G6P]/dt$ | $= v_{G6PI} - v_{PGM}$ |
| $d[G1P]/dt$ | $= -v_{starch} + v_{PGM}$ |
| $d[SBP]/dt$ | $= -v_{SBPase} + v_{ALD,SBP}$ |
| $d[S7P]/dt$ | $= v_{SBPase} - \mathbf{x_0}$ |
| $d[E4P]/dt$ | $= -v_{ALD,SBP} + \mathbf{x_0}$ |
| $d[X5P]/dt$ | $= -\mathbf{x_0} + \mathbf{x_1} + \mathbf{x_2}$ |
| $d[R5P]/dt$ | $= -\mathbf{x_0} + \mathbf{x_1}$ |
| $d[RuBP]/dt$ | $= v_{PRK} - v_{Rubisco}$ |
| $d[Ru5P]/dt$ | $= -v_{PRK} + \mathbf{x_0} + \mathbf{x_1}$ |
| $d[ATP]/dt$ | $= v_{ATPsynth}\,f_{conv} - v^{ex}_{ATP} - v_{starch} - v_{PRK} - \mathbf{x_0}$ |
| $d[Fd_{ox}]/dt$ | $= 2\,v_{NDH} + 2\,v_{FNR} - v_{\to P700^{+}F_A} - v^{(3)}_{\to P700F_A} + 2\,\mathbf{x_0} + \mathbf{x_1}$ |
| $d[LHC]/dt$ | $= -\mathbf{x_0} + \mathbf{x_1}$ |
| $d[NADPH]/dt$ | $= v_{FNR}\,f_{conv} - v^{ex}_{NADPH} - v_{GAPDH} - v_{MDAR2} - \mathbf{x_0}$ |
| $d[PC_{ox}]/dt$ | $= -2\,v_{b_6f} + v_{\to P700F_A^-} + v^{(5)}_{\to P700F_A}$ |
| $d[PQ_{ox}]/dt$ | $= -0.5\,v_{B_2\to B_0} - v_{NDH} + v_{PTOX} + v_{b_6f} - v_{ndh} - \mathbf{x_0}$ |
| $d[PsbS_{deprot}]/dt$ | $= v^{deprot}_{LHC} - v^{prot}_{LHC}$ |
| $d[Vx]/dt$ | $= -\mathbf{x_0} + \mathbf{x_1}$ |
| $d[MDA]/dt$ | $= 2\,(-v_{MDAR1} - v_{MDAR2} + \mathbf{x_0})$ |
| $d[H_2O_2]/dt$ | $= v_{Mehler,1}\,f_{conv} + v_{Mehler,2}\,f_{conv} + v^{inact,2}_{Trx}\,f_{conv} - \mathbf{x_0}$ |
| $d[DHA]/dt$ | $= v_{MDAR1} - \mathbf{x_0}$ |

*Continued on next page*

Table S2: Ordinary differential equations of the model (continued)

| Variable | Differential equation |
|---:|---|
| $\mathrm{d}[\mathrm{GSSG}]/\mathrm{d}t$ | $= \mathbf{x_0} - \mathbf{x_1}$ |
| $\mathrm{d}[\mathrm{Trx_{ox}}]/\mathrm{d}t$ | $= 5\, v_{\mathrm{Trx}}^{\mathrm{act}} + v_{\mathrm{Trx}}^{\mathrm{inact,2}} - \mathbf{x_0}$ |
| $\mathrm{d}[\mathrm{E_{inactive}}]/\mathrm{d}t$ | $= 5\,(-v_{\mathrm{Trx}}^{\mathrm{act}} + v_{\mathrm{Trx}}^{\mathrm{inact}})$ |
| $\mathrm{d}[\mathrm{P700\,F_A}]/\mathrm{d}t$ | $= -v_{\mathrm{PSI}} + 2\, v_{\mathrm{Mehler,1}} + v_{\rightarrow\mathrm{P700F_A}}^{(3)} + v_{\rightarrow\mathrm{P700F_A}}^{(5)}$ |
| $\mathrm{d}[\mathrm{P700^+F_A^-}]/\mathrm{d}t$ | $= v_{\mathrm{PSI}} - 2\, v_{\mathrm{Mehler,2}} - v_{\rightarrow\mathrm{P700^+F_A}} - v_{\rightarrow\mathrm{P700F_A^-}}$ |
| $\mathrm{d}[\mathrm{P700\,F_A^-}]/\mathrm{d}t$ | $= -2\, v_{\mathrm{Mehler,1}} + v_{\rightarrow\mathrm{P700F_A^-}} - v_{\rightarrow\mathrm{P700F_A}}^{(3)}$ |
| $\mathrm{d}[\mathrm{B_0}]/\mathrm{d}t$ | $= -v_{\mathrm{B_0\rightarrow B_1}} + v_{\mathrm{B_1\rightarrow B_0}}^{\mathrm{F}} + v_{\mathrm{B_1\rightarrow B_0}}^{\mathrm{Q}} + v_{\mathrm{B_2\rightarrow B_0}}$ |
| $\mathrm{d}[\mathrm{B_1}]/\mathrm{d}t$ | $= v_{\mathrm{B_0\rightarrow B_1}} - v_{\mathrm{B_1\rightarrow B_0}}^{\mathrm{F}} - v_{\mathrm{B_1\rightarrow B_0}}^{\mathrm{Q}} - v_{\mathrm{B_1\rightarrow B_2}}$ |
| $\mathrm{d}[\mathrm{B_2}]/\mathrm{d}t$ | $= v_{\mathrm{B_1\rightarrow B_2}} - v_{\mathrm{B_2\rightarrow B_0}} - v_{\mathrm{B_2\rightarrow B_3}} + v_{\mathrm{B_3\rightarrow B_2}}^{\mathrm{F}} + v_{\mathrm{B_3\rightarrow B_2}}^{\mathrm{Q}}$ |
| $\mathrm{dpH_{lumen}}/\mathrm{d}t$ | $= (-v_{\mathrm{B_1\rightarrow B_2}} + v_{\mathrm{ATPsynth}}\, n_{\mathrm{H^+/ATP}} + v_{\mathrm{KEA3}} - 4\, v_{\mathrm{NDH}} - 4\, v_{\mathrm{b_6f}} + v_{\mathrm{leak}})/b_{\mathrm{H}}$ |
| $\mathrm{dpH_{stroma}}/\mathrm{d}t$ | $= (v_{\mathrm{B_2\rightarrow B_0}} - v_{\mathrm{ATPsynth}}\, n_{\mathrm{H^+/ATP}} - v_{\mathrm{KEA3}} + 4\, v_{\mathrm{NDH}} + 4\, v_{\mathrm{b_6f}} - v_{\mathrm{leak}})/b_{\mathrm{H}}^{\mathrm{stroma}}$ |
| $\mathrm{d}a_{\mathrm{ATPsynth}}/\mathrm{d}t$ | $= v_{\mathrm{ATPsynth}}^{\mathrm{act}}$ |
| $\mathrm{d}\Delta\psi/\mathrm{d}t$ | $= V_{\mathrm{charge}}\,(v_{\mathrm{B_0\rightarrow B_1}} - v_{\mathrm{B_1\rightarrow B_0}}^{\mathrm{F}} - v_{\mathrm{B_1\rightarrow B_0}}^{\mathrm{Q}} - v_{\mathrm{ClCe}} + v_{\mathrm{Cl\,leak}} - v_{\mathrm{ATPsynth}}\, n_{\mathrm{H^+/ATP}} +$ <br> $4\, v_{\mathrm{NDH}} + v_{\mathrm{PSI}} - v_{\mathrm{VCCN1}} + 2\, v_{\mathrm{b_6f}} - v_{\mathrm{leak}} - v_{\mathrm{K^+\,channel}})$ |
| $\mathrm{d}[\mathrm{K^+_{stroma}}]/\mathrm{d}t$ | $= 0.032\,(-v_{\mathrm{KEA3}} + v_{\mathrm{K^+\,channel}})$ |
| $\mathrm{d}[\mathrm{Cl^-_{stroma}}]/\mathrm{d}t$ | $= 0.032\,(-v_{\mathrm{ClCe}} + v_{\mathrm{Cl\,leak}} - v_{\mathrm{VCCN1}})$ |
| $\mathrm{d}[\mathrm{K^+_{lumen}}]/\mathrm{d}t$ | $= 0.25\,(v_{\mathrm{KEA3}} - v_{\mathrm{K^+\,channel}})$ |
| $\mathrm{d}[\mathrm{Cl^-_{lumen}}]/\mathrm{d}t$ | $= 0.25\,(v_{\mathrm{ClCe}} - v_{\mathrm{Cl\,leak}} + v_{\mathrm{VCCN1}})$ |

#### 1.3 Rate equations

Table S3: Rate equations of the model reactions. Square brackets denote concentrations.

| Reaction | Rate equation |
|---|---|
| $v_{\mathrm{FTR}}$ | $= [\mathrm{Fd_{red}}]\,[\mathrm{Trx_{ox}}]\,k_{\mathrm{FTR}}$ |
| $v^{\mathrm{act}}_{\mathrm{Trx}}$ | $= [\mathrm{E_{inactive}}]\,[\mathrm{Trx_{red}}]\,k^{\mathrm{f}}_{\mathrm{TrxActivation}}$ |
| $v^{\mathrm{inact}}_{\mathrm{Trx}}$ | $= [\mathrm{E_{active}}]\,k^{\mathrm{f}}_{\mathrm{TrxInactivation}}$ |
| $v_{\mathrm{ATPsynth}}$ | $= f^{\mathrm{pmf}}_{\mathrm{ATPsynth}}\,\mathrm{a_{ATPsynth}}\,k_{\mathrm{ATPsynthase}}\,([\mathrm{ADP}]/f_{\mathrm{conv}} - \frac{[\mathrm{ATP}]}{f_{\mathrm{conv}}\,K^{\mathrm{eq}}_{\mathrm{ATPsynth}}})$ |
| $v^{\mathrm{prot}}_{\mathrm{LHC}}$ | $= \frac{[\mathrm{PsbS_{deprot}}]\,k^{\mathrm{prot}}_{\mathrm{LHC}}\,[\mathrm{H^+_{lumen}}]^{n^{\mathrm{LHC}}_{\mathrm{H}}}}{\mathcal{D}}$ <br> $\mathcal{D} = [\mathrm{H^+_{lumen}}]^{n^{\mathrm{LHC}}_{\mathrm{H}}} + (4000 \cdot 10^{-k^{\mathrm{sat}}_{\mathrm{LHC\,protonation}}})^{n^{\mathrm{LHC}}_{\mathrm{H}}}$ |
| $v^{\mathrm{deprot}}_{\mathrm{LHC}}$ | $= [\mathrm{PsbS_{prot}}]\,k^{\mathrm{deprot}}_{\mathrm{LHC}}$ |
| $v_{\mathrm{CEF}}$ | $= [\mathrm{Fd_{red}}]^2\,[\mathrm{PQ_{ox}}]\,k_{\mathrm{CEF}}$ |
| $v_{\mathrm{VDE}}$ | $= \frac{[\mathrm{Vx}]\,k_{\mathrm{VDE}}\,[\mathrm{H^+_{lumen}}]^{n^{\mathrm{VDE}}_{\mathrm{H}}}}{[\mathrm{H^+_{lumen}}]^{n^{\mathrm{VDE}}_{\mathrm{H}}} + (4000 \cdot 10^{-k^{\mathrm{sat}}_{\mathrm{VDE}}})^{n^{\mathrm{VDE}}_{\mathrm{H}}}}$ |
| $v_{\mathrm{ZE}}$ | $= [\mathrm{Zx}]\,k_{\mathrm{ZE}}$ |
| $v_{\mathrm{FNR}}$ | $= \frac{\mathcal{N}}{\mathcal{D}}$ <br> $\mathcal{N} = V^{\mathrm{max}}_{\mathrm{FNR}}\,(\frac{[\mathrm{NADP^+}]\,(\frac{[\mathrm{Fd_{red}}]}{K^{\mathrm{FNR}}_{m,\mathrm{Fd_{red}}}})^2}{f_{\mathrm{conv}}\,K^{\mathrm{FNR}}_{m,\mathrm{NADP^+}}} - \frac{[\mathrm{NADPH}]\,(\frac{[\mathrm{Fd_{ox}}]}{K^{\mathrm{FNR}}_{m,\mathrm{Fd_{red}}}})^2}{f_{\mathrm{conv}}\,K^{\mathrm{eq}}_{\mathrm{FNR}}\,K^{\mathrm{FNR}}_{m,\mathrm{NADP^+}}})$ <br> $\mathcal{D} = (\frac{[\mathrm{NADP^+}]}{f_{\mathrm{conv}}\,K^{\mathrm{FNR}}_{m,\mathrm{NADP^+}}} + 1)\,(\frac{[\mathrm{Fd_{red}}]}{K^{\mathrm{FNR}}_{m,\mathrm{Fd_{red}}}} + (\frac{[\mathrm{Fd_{red}}]}{K^{\mathrm{FNR}}_{m,\mathrm{Fd_{red}}}})^2 + 1) +$ <br> $(\frac{[\mathrm{NADPH}]}{f_{\mathrm{conv}}\,K^{\mathrm{FNR}}_{m,\mathrm{NADP^+}}} + 1)\,(\frac{[\mathrm{Fd_{ox}}]}{K^{\mathrm{FNR}}_{m,\mathrm{Fd_{red}}}} + (\frac{[\mathrm{Fd_{ox}}]}{K^{\mathrm{FNR}}_{m,\mathrm{Fd_{red}}}})^2 + 1) - 1$ |
| $v_{\mathrm{ndh}}$ | $= [\mathrm{PQ_{ox}}]\,k_{\mathrm{ndh}}$ |
| $v_{\mathrm{leak}}$ | $= k_{\mathrm{leak}}\,(\mathrm{pH_{stroma}} - \mathrm{pH_{lumen}})$ |
| $v_{\mathrm{PTOX}}$ | $= [\mathrm{O_2}]_{\mathrm{lumen}}\,[\mathrm{PQ_{red}}]\,k_{\mathrm{PTOX}}$ |
| $v^{1\to2}_{\mathrm{ST}}$ | $= \frac{[\mathrm{LHC}]\,k_{\mathrm{STN7}}}{(\frac{[\mathrm{PQ_{ox}}]}{\mathrm{PQ_{tot}}\,K^{\mathrm{ST}}_{m}})^{n_{\mathrm{ST}}} + 1}$ |
| $v^{2\to1}_{\mathrm{ST}}$ | $= [\mathrm{LHC_{prot}}]\,k_{\mathrm{PPH1}}$ |

*Continued on next page*

Table S3: Rate equations of the model reactions (continued)

| Reaction | Rate equation |
|---|---|
| $v_{\text{Rubisco}}$ | $= \frac{[\text{CO}_2]\,[\text{RuBP}]\,V^{\max}_{\text{Rubisco}}}{\mathcal{D}}$ <br> $\mathcal{D} = ([\text{CO}_2] + K^{\text{Rubisco}}_{m,\text{CO}_2})\,([\text{RuBP}] + K^{\text{Rubisco}}_{m,\text{RuBP}}\,(\frac{[\text{3PGA}]}{K^{\text{Rubisco}}_{i,\text{3PGA}}} + \frac{[\text{FBP}]}{K^{\text{Rubisco}}_{i,\text{FBP}}} +$ <br> $\frac{[\text{NADPH}]}{K^{\text{Rubisco}}_{i,\text{NADPH}}} + \frac{[\text{P}_\text{i}]}{K^{\text{Rubisco}}_{i,\text{P}_\text{i}}} + \frac{[\text{SBP}]}{K^{\text{Rubisco}}_{i,\text{SBP}}} + 1))$ |
| $v_{\text{PGK}}$ | $= k^{\text{re}}_{\text{PGK}}\,([\text{3PGA}]\,[\text{ATP}] - [\text{ADP}]\,[\text{BPGA}]/K^{\text{eq}}_{\text{PGK}})$ |
| $v_{\text{GAPDH}}$ | $= k^{\text{re}}_{\text{GAPDH}}\,([\text{BPGA}]\,[\text{NADPH}]\,[\text{H}^+_{\text{stroma}}] -$ <br> $[\text{GAP}]\,[\text{NADP}^+]\,[\text{P}_\text{i}]/K^{\text{eq}}_{\text{GAPDH}})$ |
| $v_{\text{TPI}}$ | $= k^{\text{re}}_{\text{TPI}}\,(-\frac{[\text{DHAP}]}{K^{\text{eq}}_{\text{TPI}}} + [\text{GAP}])$ |
| $v_{\text{ALD,FBP}}$ | $= k^{\text{re}}_{\text{ALD,FBP}}\,([\text{DHAP}]\,[\text{GAP}] - \frac{[\text{FBP}]}{K^{\text{eq}}_{\text{ALD,FBP}}})$ |
| $v_{\text{ALD,SBP}}$ | $= k^{\text{re}}_{\text{ALD,SBP}}\,([\text{DHAP}]\,[\text{E4P}] - \frac{[\text{SBP}]}{K^{\text{eq}}_{\text{ALD,SBP}}})$ |
| $v_{\text{FBPase}}$ | $= \frac{[\text{FBP}]\,V^{\max}_{\text{FBPase}}}{[\text{FBP}] + K^{\text{FBPase}}_{m,\text{S}}\,([\text{F6P}]/K^{\text{FBPase}}_{i,\text{F6P}} + [\text{P}_\text{i}]/K^{\text{FBPase}}_{i,\text{P}_\text{i}} + 1)}$ |
| $v_{\text{TK,F6P}}$ | $= k^{\text{re}}_{\text{TK,F6P}}\,(-\frac{[\text{E4P}]\,[\text{X5P}]}{K^{\text{eq}}_{\text{TK,F6P}}} + [\text{F6P}]\,[\text{GAP}])$ |
| $v_{\text{TK,S7P}}$ | $= k^{\text{re}}_{\text{TK,S7P}}\,([\text{GAP}]\,[\text{S7P}] - \frac{[\text{R5P}]\,[\text{X5P}]}{K^{\text{eq}}_{\text{TK,S7P}}})$ |
| $v_{\text{SBPase}}$ | $= \frac{[\text{SBP}]\,V^{\max}_{\text{SBPase}}}{[\text{SBP}] + K^{\text{SBPase}}_{m,\text{S}}\,([\text{P}_\text{i}]/K^{\text{SBPase}}_{i,\text{P}_\text{i}} + 1)}$ |
| $v_{\text{RPI}}$ | $= k^{\text{re}}_{\text{RPI}}\,([\text{R5P}] - \frac{[\text{Ru5P}]}{K^{\text{eq}}_{\text{RPI}}})$ |
| $v_{\text{RPE}}$ | $= k^{\text{re}}_{\text{RPE}}\,(-\frac{[\text{Ru5P}]}{K^{\text{eq}}_{\text{RPE}}} + [\text{X5P}])$ |
| $v_{\text{PRK}}$ | $= \frac{[\text{ATP}]\,[\text{Ru5P}]\,V^{\max}_{\text{PRK}}}{\mathcal{D}}$ <br> $\mathcal{D} = ([\text{Ru5P}] + K^{\text{PRK}}_{m,\text{Ru5P}}\,([\text{3PGA}]/K^{\text{PRK}}_{i,\text{3PGA}} + [\text{P}_\text{i}]/K^{\text{PRK}}_{i,\text{P}_\text{i}} +$ <br> $[\text{RuBP}]/K^{\text{PRK}}_{i,\text{RuBP}} + 1))\,([\text{ATP}]\,([\text{ADP}]/K^{\text{PRK}}_{i,4} + 1) +$ <br> $K^{\text{PRK}}_{m,\text{ATP}}\,([\text{ADP}]/K^{\text{PRK}}_{i,5} + 1))$ |
| $v_{\text{G6PI}}$ | $= k^{\text{re}}_{\text{G6PI}}\,([\text{F6P}] - [\text{G6P}]/K^{\text{eq}}_{\text{G6PI}})$ |
| $v_{\text{PGM}}$ | $= k^{\text{re}}_{\text{PGM}}\,(-[\text{G1P}]/K^{\text{eq}}_{\text{PGM}} + [\text{G6P}])$ |
| $v^{\text{ex}}_{\text{3PGA}}$ | $= \frac{[\text{3PGA}]\,V^{\max}_{\text{TPT}}}{N_{\text{TPT}}\,K^{\text{export}}_{m,\text{3PGA}}}$ |
| $v^{\text{ex}}_{\text{GAP}}$ | $= \frac{[\text{GAP}]\,V^{\max}_{\text{TPT}}}{N_{\text{TPT}}\,K^{\text{export}}_{m,\text{GAP}}}$ |
| $v^{\text{ex}}_{\text{DHAP}}$ | $= \frac{[\text{DHAP}]\,V^{\max}_{\text{TPT}}}{N_{\text{TPT}}\,K^{\text{export}}_{m,\text{DHAP}}}$ |

*Continued on next page*

Table S3: Rate equations of the model reactions (continued)

| Reaction | Rate equation |
|---|---|
| $v_{\mathrm{starch}}$ | $= \frac{[\mathrm{ATP}]\,[\mathrm{G1P}]\,V^{\mathrm{max}}_{\mathrm{starch}}}{\mathcal{D}}$ <br> $\mathcal{D} = ([\mathrm{G1P}] + K^{\mathrm{starch}}_{m,\mathrm{G1P}})\,(\frac{[\mathrm{P_i}]\,K^{\mathrm{starch}}_{m,\mathrm{ATP}}}{[\mathrm{3PGA}]\,K^{\mathrm{starch}}_{i,\mathrm{3PGA}}+[\mathrm{F6P}]\,K^{\mathrm{starch}}_{i,\mathrm{F6P}}+[\mathrm{FBP}]\,K^{\mathrm{starch}}_{i,\mathrm{FBP}}} +$ <br> $([\mathrm{ATP}] + K^{\mathrm{starch}}_{m,\mathrm{ATP}})\,([\mathrm{ADP}]/K^{\mathrm{starch}}_{i} + 1))$ |
| $v_{\mathrm{MDAR1}}$ | $= [\mathrm{MDA}]^2\,k_{\mathrm{MDAR1}}$ |
| $v_{\mathrm{MDAR2}}$ | $= \frac{[\mathrm{MDA}]\,[\mathrm{NADPH}]\,V^{\mathrm{max}}_{\mathrm{MDAR2}}}{\mathcal{D}}$ <br> $\mathcal{D} = [\mathrm{MDA}]\,[\mathrm{NADPH}] + [\mathrm{MDA}]\,K^{\mathrm{MDAR2}}_{m,\mathrm{NADPH}} + [\mathrm{NADPH}]\,K^{\mathrm{MDAR2}}_{m,\mathrm{MDA}} +$ <br> $K^{\mathrm{MDAR2}}_{m,\mathrm{MDA}}\,K^{\mathrm{MDAR2}}_{m,\mathrm{NADPH}}$ |
| $v_{\mathrm{APX}}$ | $= \frac{[\mathrm{H_2O_2}]\,X_{\mathrm{T}}\,[\mathrm{ASC}]}{\mathcal{D}}$ <br> $\mathcal{D} = [\mathrm{H_2O_2}]\,[\mathrm{ASC}]\,(1/k^{\mathrm{f}}_5 + 1/k^{\mathrm{f}}_3) + [\mathrm{H_2O_2}]/k^{\mathrm{f}}_4 + \frac{[\mathrm{H_2O_2}]\,k^{\mathrm{r}}_4}{k^{\mathrm{f}}_4\,k^{\mathrm{f}}_5} +$ <br> $[\mathrm{H_2O_2}]/k^{\mathrm{f}}_2 + \frac{[\mathrm{H_2O_2}]\,k^{\mathrm{r}}_2}{k^{\mathrm{f}}_2\,k^{\mathrm{f}}_3} + [\mathrm{ASC}]/k^{\mathrm{f}}_1 + \frac{k^{\mathrm{r}}_1}{k^{\mathrm{f}}_1\,k^{\mathrm{f}}_2} + \frac{k^{\mathrm{r}}_1\,k^{\mathrm{r}}_2}{k^{\mathrm{f}}_1\,k^{\mathrm{f}}_2\,k^{\mathrm{f}}_3}$ |
| $v_{\mathrm{GR}}$ | $= \frac{[\mathrm{GSSG}]\,[\mathrm{NADPH}]\,V^{\mathrm{max}}_{\mathrm{GR}}}{\mathcal{D}}$ <br> $\mathcal{D} = [\mathrm{GSSG}]\,[\mathrm{NADPH}] + [\mathrm{GSSG}]\,K^{\mathrm{GR}}_{m,\mathrm{NADPH}} +$ <br> $[\mathrm{NADPH}]\,K^{\mathrm{GR}}_{m,\mathrm{GSSG}} + K^{\mathrm{GR}}_{m,\mathrm{GSSG}}\,K^{\mathrm{GR}}_{m,\mathrm{NADPH}}$ |
| $v_{\mathrm{DHAR}}$ | $= \frac{[\mathrm{DHA}]\,[\mathrm{GSH}]\,V^{\mathrm{max}}_{\mathrm{DHAR}}}{[\mathrm{DHA}]\,[\mathrm{GSH}] + [\mathrm{DHA}]\,K^{\mathrm{DHAR}}_{m,\mathrm{GSH}} + [\mathrm{GSH}]\,K^{\mathrm{DHAR}}_{m,\mathrm{DHA}} + K_{\mathrm{DHAR}}}$ |
| $v^{\mathrm{ex}}_{\mathrm{ATP}}$ | $= [\mathrm{ATP}]\,k^{\mathrm{ex}}_{\mathrm{ATP}}$ |
| $v^{\mathrm{ex}}_{\mathrm{NADPH}}$ | $= [\mathrm{NADPH}]\,k^{\mathrm{ex}}_{\mathrm{NADPH}}$ |
| $v_{\to\mathrm{P700F^-_A}}$ | $= [\mathrm{P700^+F^-_A}]\,[\mathrm{PC_{red}}]\,k_{\mathrm{PC,ox}} - [\mathrm{P700\,F^-_A}]\,[\mathrm{PC_{ox}}]\,k_{\mathrm{PC,ox}}/K^{\mathrm{eq}}_{\mathrm{PC,P700}}$ |
| $v^{(3)}_{\to\mathrm{P700F_A}}$ | $= [\mathrm{Fd_{ox}}]\,[\mathrm{P700\,F^-_A}]\,k_{\mathrm{Fd,red}} - [\mathrm{Fd_{red}}]\,[\mathrm{P700\,F_A}]\,k_{\mathrm{Fd,red}}/K^{\mathrm{eq}}_{\mathrm{F_A,Fd}}$ |
| $v_{\to\mathrm{P700^+F_A}}$ | $= [\mathrm{Fd_{ox}}]\,[\mathrm{P700^+F^-_A}]\,k_{\mathrm{Fd,red}} - [\mathrm{Fd_{red}}]\,[\mathrm{P700^+F_A}]\,k_{\mathrm{Fd,red}}/K^{\mathrm{eq}}_{\mathrm{F_A,Fd}}$ |
| $v^{(5)}_{\to\mathrm{P700F_A}}$ | $= [\mathrm{P700^+F_A}]\,[\mathrm{PC_{red}}]\,k_{\mathrm{PC,ox}} - [\mathrm{P700\,F_A}]\,[\mathrm{PC_{ox}}]\,k_{\mathrm{PC,ox}}/K^{\mathrm{eq}}_{\mathrm{PC,P700}}$ |
| $v_{\mathrm{PSI}}$ | $= [\mathrm{P700\,F_A}]\,\mathrm{PPFD}\,(1 - \sigma_{\mathrm{PSII}})$ |
| $v_{\mathrm{Mehler,1}}$ | $= [\mathrm{O_2}]_{\mathrm{lumen}}\,[\mathrm{P700\,F^-_A}]\,k_{\mathrm{Mehler}}$ |
| $v_{\mathrm{Mehler,2}}$ | $= [\mathrm{O_2}]_{\mathrm{lumen}}\,[\mathrm{P700^+F^-_A}]\,k_{\mathrm{Mehler}}$ |
| $v_{\mathrm{B_0\to B_1}}$ | $= [\mathrm{B_0}]\,\mathrm{PPFD}\,\sigma_{\mathrm{PSII}}$ |
| $v^{\mathrm{Q}}_{\mathrm{B_1\to B_0}}$ | $= [\mathrm{B_1}]\,(Q\,k^{\mathrm{Qslope}}_{\mathrm{H}} + k_{\mathrm{H0}})$ |

*Continued on next page*

Table S3: Rate equations of the model reactions (continued)

| Reaction | Rate equation |
|---|---|
| $v^{\mathrm{F}}_{\mathrm{B_1 \to B_0}}$ | $= [\mathrm{B_1}]\, k_{\mathrm{F}}$ |
| $v_{\mathrm{B_1 \to B_2}}$ | $= [\mathrm{B_1}]\, k_2$ |
| $v_{\mathrm{B_2 \to B_0}}$ | $= -[\mathrm{B_0}]\, [\mathrm{PQ_{red}}]\, k_{\mathrm{PQ,red}} / K^{\mathrm{eq}}_{\mathrm{PQ}} + [\mathrm{B_2}]\, [\mathrm{PQ_{ox}}]\, k_{\mathrm{PQ,red}}$ |
| $v_{\mathrm{B_2 \to B_3}}$ | $= [\mathrm{B_2}]\, \mathrm{PPFD}\, \sigma_{\mathrm{PSII}}$ |
| $v^{\mathrm{F}}_{\mathrm{B_3 \to B_2}}$ | $= [\mathrm{B_3}]\, k_{\mathrm{F}}$ |
| $v^{\mathrm{Q}}_{\mathrm{B_3 \to B_2}}$ | $= [\mathrm{B_3}]\, (Q\, k_{\mathrm{H}}^{\mathrm{Qslope}} + k_{\mathrm{H0}})$ |
| $v_{\mathrm{b_6 f}}$ | $= k^{\mathrm{dyn}}_{\mathrm{b_6 f}}\, ([\mathrm{PC_{ox}}]^2\, [\mathrm{PQ_{red}}] - [\mathrm{PC_{red}}]^2\, [\mathrm{PQ_{ox}}] / K^{\mathrm{eq}}_{\mathrm{b_6 f}})$ |
| $v^{\mathrm{act}}_{\mathrm{ATPsynth}}$ | $= -\dfrac{\mathrm{a_{ATPsynth}}\, k^{\mathrm{deactivate}}_{\mathrm{ATPsynthase}}}{3.72 \cdot 10^{-44}\, e^{20\, \mathrm{PPFD}} + 1} + \dfrac{k^{\mathrm{activate}}_{\mathrm{ATPsynthase}}\, (1 - \mathrm{a_{ATPsynth}})}{1 + 2.68 \cdot 10^{43}\, e^{-20\, \mathrm{PPFD}}}$ |
| $v_{\mathrm{KEA3}}$ | $= \max(0, a_{\mathrm{KEA3}}\, k_{\mathrm{KEA3}}\, (-[\mathrm{K^+_{lumen}}]\, \mathrm{pH_{stroma}} / [\mathrm{K^+}]_{\mathrm{tot}} +$ $[\mathrm{K^+_{stroma}}]\, \mathrm{pH_{lumen}} / [\mathrm{K^+}]_{\mathrm{tot}}))$ |
| $v_{\mathrm{K^+\,channel}}$ | $= \dfrac{0.9\, [\mathrm{K^+_{lumen}}]\, \Delta G_{\mathrm{K^+}}\, perm_{\mathrm{K^+}}}{\mathcal{D}}$ $\mathcal{D} = [\mathrm{K^+_{stroma}}]\, (e^{1000\, \mathrm{K}^{+\Delta\psi}_{\mathrm{threshold}} - 1000\, \Delta\psi} + 1)$ |
| $v_{\mathrm{VCCN1}}$ | $= \dfrac{0.9\, \Delta G_{\mathrm{Cl^-}}\, [\mathrm{Cl^-_{stroma}}]\, k_{\mathrm{VCCN1}}}{\mathcal{D}}$ $\mathcal{D} = [\mathrm{Cl^-_{lumen}}]\, (e^{1000\, \mathrm{VCCN1}^{\Delta\psi}_{\mathrm{threshold}} - 1000\, \Delta\psi} + 1)$ |
| $v_{\mathrm{Cl\,leak}}$ | $= \dfrac{0.9\, k_{\mathrm{Cl\,leak}}\, ([\mathrm{Cl^-_{lumen}}] - [\mathrm{Cl^-_{stroma}}])^2}{\mathrm{Cl_{tot,2}}\, (e^{10\, \mathrm{PQ^{Cl\,leak}_{thr}} - 10\, [\mathrm{PQ_{ox}}]} + 1)}$ |
| $v_{\mathrm{NDH}}$ | $= \dfrac{0.9 \cdot 10^{\mathrm{pH_{lumen}} - 6.5}\, [\mathrm{Fd_{red}}]^2\, [\mathrm{PQ_{ox}}]\, k_{\mathrm{NDH}}}{\mathcal{D}}$ $\mathcal{D} = (1 + 7.38\, e^{-100\, [\mathrm{P700^+ F_A^-}]})\, (10^{\mathrm{pH_{lumen}} - 6.5} + 0.5)$ |
| $v_{\mathrm{ClCe}}$ | $= a_{\mathrm{ClCe}}\, k_{\mathrm{ClCe}}\, (-[\mathrm{Cl^-_{lumen}}] + [\mathrm{Cl^-_{stroma}}])$ |
| $v^{\mathrm{inact,2}}_{\mathrm{Trx}}$ | $= [\mathrm{Trx_{red}}]\, k2^{\mathrm{Trx}}_{\mathrm{inactivation}}$ |

##### 1.4 Parameters

Table S4: All model parameters. Fitted values are given at the final optimum; "Inherited" denotes values taken from Saadat et al. (2021).

| Parameter | Value | Parameter category |
|---|---:|---|
| **Environment and physical constants** | | |
| $b_{\mathrm{H}}$ | 200.00 | Assumed |
| $[\mathrm{CO_2}]$ | 0.0136 | Literature-derived; atmospheric $CO_2$ condition |
| $f_{\mathrm{conv}}$ | 0.0320 | Inherited |
| $F$ | 96.48 | Inherited |
| $[\mathrm{O_2}]_{\mathrm{lumen}}$ | 8.00 | Inherited |
| PPFD | 100.00 | Inherited |
| $R$ | 0.0083 | Inherited |
| $b_{\mathrm{H}}^{\mathrm{stroma}}$ | 500.00 | Literature-derived / assumed; larger buffering than in the lumen, chosen so that stromal pH stays between 7 and 8 |
| $T$ | 298.00 | Inherited |
| $V_{\mathrm{charge}}$ | 0.00077 | Assumed; keeps the membrane potential in the reported physiological range |
| **Ion transport and pmf partitioning** | | |
| $\mathrm{PQ}_{\mathrm{thr}}^{\mathrm{Cl\,leak}}$ | 6.61 | Fitted; bounds based on assumed activity of an unknown channel |
| $[\mathrm{Cl^-}]_{\mathrm{tot}}$ | 15.65 | Fitted |
| $\mathrm{ClCe}_{\mathrm{threshold}}^{\mathrm{ATP}}$ | 0.0358 | Fitted |
| $k_{\mathrm{Cl\,leak}}$ | 3.28 | Fitted |
| $k_{\mathrm{ClCe}}$ | 1.00 | Fitted |
| $\mathrm{K}_{\mathrm{threshold}}^{+\Delta\psi}$ | 0.0526 | Fitted |
| $k_{\mathrm{KEA3}}$ | 17.86 | Fitted |
| $[\mathrm{K^+}]_{\mathrm{tot}}$ | 68.97 | Fitted |
| $k_{\mathrm{VCCN1}}$ | 3.01 | Fitted |
| $\mathrm{KEA3}_{\mathrm{threshold}}^{\mathrm{ATP}}$ | 0.01 | Assumed / literature-derived, based on reported activity of KEA3 |
| $\mathrm{KEA3}_{\mathrm{pHreg}}^{\mathrm{KEA3}}$ | 7.62 | Fitted; bounds based on reported activity of KEA3 |
| $perm_{\mathrm{K+}}$ | 5.69 | Fitted |
| $\mathrm{VCCN1}_{\mathrm{threshold}}^{\Delta\psi}$ | 0.0044 | Fitted |
| **ATP synthase** | | |
| $\Delta G_{\mathrm{ATP}}^{0}$ | 30.60 | Inherited |
| $n_{\mathrm{H+/ATP}}$ | 4.67 | Inherited |
| $k_{\mathrm{ATPsynthase}}^{\mathrm{activate}}$ | 0.0012 | Fitted |
| $k_{\mathrm{ATPsynthase}}^{\mathrm{deactivate}}$ | 0.00010 | Fitted |

*Continued on next page*

Table S4: All model parameters (continued)

| Parameter | Value | Parameter category |
|---|---|---|
| $k_{\mathrm{ATPsynthase}}$ | 20.00 | Assumed |
| $[\mathrm{P_i}]_{\mathrm{mol}}$ | 0.01 | Inherited |
| **Photosynthetic electron transport** | | |
| $\mathrm{AP_{tot}}$ | 2.55 | Inherited |
| $\mathrm{b_6f_{tot}}$ | 1.00 | Literature-derived |
| $E_{0,\mathrm{FNR}}$ | 3.00 | Inherited |
| $E^0_{\mathrm{FA}}$ | -0.55 | Inherited |
| $E^0_{\mathrm{Fd}}$ | -0.43 | Inherited |
| $E^0_{\mathrm{NADP}}$ | -0.11 | Inherited |
| $E^0_{\mathrm{P700}}$ | 0.48 | Inherited |
| $E^0_{\mathrm{PC}}$ | 0.38 | Inherited |
| $E^0_{\mathrm{PQ}}$ | 0.35 | Inherited |
| $E^0_{\mathrm{QA}}$ | -0.14 | Inherited |
| $\mathrm{Fd_{tot}}$ | 5.00 | Inherited |
| $k_2$ | $5.00 \times 10^9$ | Inherited |
| $k_{\mathrm{NDH}}$ | 6.97 | Fitted |
| $k^{\mathrm{cat}}_{\mathrm{FNR}}$ | 500.00 | Inherited |
| $k_{\mathrm{F}}$ | $6.25 \times 10^8$ | Inherited |
| $k_{\mathrm{CEF}}$ | 1.00 | Inherited |
| $k_{\mathrm{ndh}}$ | 0.0020 | Inherited |
| $k_{\mathrm{leak}}$ | 10.00 | Inherited |
| $k_{\mathrm{Fd,red}}$ | $2.50 \times 10^5$ | Inherited |
| $k_{\mathrm{H}}$ | $5.00 \times 10^9$ | Inherited |
| $k_{\mathrm{H0}}$ | $5.00 \times 10^8$ | Inherited |
| $k^{\mathrm{Qslope}}_{\mathrm{H}}$ | $5.00 \times 10^9$ | Added with dynamic photosystems / literature-derived |
| $K^{\mathrm{FNR}}_{m,\mathrm{Fd_{red}}}$ | 1.56 | Inherited |
| $K^{\mathrm{FNR}}_{m,\mathrm{NADP^+}}$ | 0.22 | Inherited |
| $K^{\mathrm{ST}}_{m}$ | 0.20 | Inherited |
| $k_{\mathrm{Mehler}}$ | 1.00 | Inherited |
| $k_{\mathrm{PC,ox}}$ | 2500.00 | Inherited |
| $k_{\mathrm{PPH1}}$ | 0.0013 | Inherited |
| $k_{\mathrm{PQ,red}}$ | 250.00 | Inherited |
| $k_{\mathrm{PTOX}}$ | 0.01 | Inherited |
| $k_{\mathrm{STN7}}$ | 0.0035 | Inherited |
| $\mathrm{LHC_{tot}}$ | 1.00 | Inherited |
| $v^{\mathrm{max}}_{\mathrm{b_6f}}$ | 300.00 | Literature-derived / assumed |
| $n_{\mathrm{ST}}$ | 2.00 | Inherited |
| $\mathrm{NADP_{tot}}$ | 0.80 | Inherited |

*Continued on next page*

Table S4: All model parameters (continued)

| Parameter | Value | Parameter category |
|---|---:|---|
| $n_{\mathrm{H}}^{\mathrm{b_6f}}$ | 3.00 | Assumed |
| $\mathrm{PC_{tot}}$ | 4.00 | Inherited |
| $\mathrm{p}K_{\mathrm{reg}}$ | 6.30 | Fitted |
| $\mathrm{PQ_{tot}}$ | 17.50 | Inherited |
| $\psi_{\mathrm{reg}}$ | 3.00 | – |
| $\mathrm{PSI_{tot}}$ | 2.50 | Inherited |
| $\mathrm{PSII_{tot}}$ | 2.50 | Inherited |
| $\sigma_{\mathrm{PSI}}^{\mathrm{static}}$ | 0.37 | Inherited |
| $\sigma_{\mathrm{PSII}}^{\mathrm{static}}$ | 0.10 | Inherited |
| **Non-photochemical quenching** | | |
| $\mathrm{Car_{tot}}$ | 1.00 | Inherited |
| $\gamma_0$ | 0.067 | Fitted |
| $\gamma_1$ | 0.94 | Fitted |
| $\gamma_2$ | 3.94 | Fitted |
| $\gamma_3$ | 3.41 | Fitted |
| $k_{\mathrm{LHC}}^{\mathrm{deprot}}$ | 0.0756 | Fitted |
| $k_{\mathrm{LHC}}^{\mathrm{prot}}$ | 0.33 | Fitted |
| $k_{\mathrm{VDE}}$ | 0.00013 | Fitted |
| $k_{\mathrm{ZE}}$ | 0.00060 | Fitted |
| $n_{\mathrm{H}}^{\mathrm{LHC}}$ | 2.16 | Fitted |
| $n_{\mathrm{H}}^{\mathrm{VDE}}$ | 19.89 | Fitted |
| $k_{\mathrm{LHC\,protonation}}^{\mathrm{sat}}$ | 5.50 | Fitted |
| $k_{\mathrm{VDE}}^{\mathrm{sat}}$ | 6.40 | Fitted |
| $\mathrm{PsbS_{tot}}$ | 1.00 | Inherited |
| **Thioredoxin system** | | |
| $E_{\mathrm{tot}}$ | 6.00 | Inherited |
| $k2_{\mathrm{inactivation}}^{\mathrm{Trx}}$ | 0.0238 | Fitted |
| $k_{\mathrm{FTR}}$ | 0.13 | Fitted |
| $k_{\mathrm{TrxActivation}}^{\mathrm{f}}$ | 0.0037 | Fitted |
| $k_{\mathrm{TrxInactivation}}^{\mathrm{f}}$ | 0.0118 | Fitted |
| $\mathrm{Trx_{tot}}$ | 1.00 | Inherited |
| **Calvin–Benson–Bassham cycle and export** | | |
| $E_{0,\mathrm{starch}}$ | 1.00 | Inherited |
| $E_{0,\mathrm{FBPase}}$ | 1.00 | Inherited |
| $E_{0,\mathrm{TPT}}$ | 1.00 | Inherited |
| $E_{0,\mathrm{PRK}}$ | 1.00 | Inherited |
| $E_{0,\mathrm{Rubisco}}$ | 1.00 | Inherited |

*Continued on next page*

Table S4: All model parameters (continued)

| Parameter | Value | Parameter category |
|---|---:|---|
| $E_{0,\mathrm{SBPase}}$ | 1.00 | Inherited |
| $k^{\mathrm{cat}}_{\mathrm{starch}}$ | 0.32 | Inherited |
| $k^{\mathrm{cat}}_{\mathrm{FBPase}}$ | 1.60 | Inherited |
| $k^{\mathrm{cat}}_{\mathrm{TPT}}$ | 2.00 | Inherited |
| $k^{\mathrm{cat}}_{\mathrm{PRK}}$ | 8.00 | Inherited |
| $k^{\mathrm{cat}}_{\mathrm{Rubisco}}$ | 2.72 | Inherited |
| $k^{\mathrm{cat}}_{\mathrm{SBPase}}$ | 0.32 | Inherited |
| $K^{\mathrm{eq}}_{\mathrm{ALD,SBP}}$ | 13.00 | Inherited |
| $K^{\mathrm{eq}}_{\mathrm{ALD,FBP}}$ | 7.10 | Inherited |
| $K^{\mathrm{eq}}_{\mathrm{G6PI}}$ | 2.30 | Inherited |
| $K^{\mathrm{eq}}_{\mathrm{GAPDH}}$ | $1.60 \times 10^{7}$ | Inherited |
| $K^{\mathrm{eq}}_{\mathrm{PGM}}$ | 0.058 | Inherited |
| $K^{\mathrm{eq}}_{\mathrm{PGK}}$ | 0.00031 | Inherited |
| $K^{\mathrm{eq}}_{\mathrm{RPI}}$ | 0.40 | Inherited |
| $K^{\mathrm{eq}}_{\mathrm{RPE}}$ | 0.67 | Inherited |
| $K^{\mathrm{eq}}_{\mathrm{TK,F6P}}$ | 0.084 | Inherited |
| $K^{\mathrm{eq}}_{\mathrm{TK,S7P}}$ | 0.85 | Inherited |
| $K^{\mathrm{eq}}_{\mathrm{TPI}}$ | 22.00 | Inherited |
| $k^{\mathrm{ex}}_{\mathrm{ATP}}$ | 0.20 | Inherited |
| $k^{\mathrm{ex}}_{\mathrm{NADPH}}$ | 0.20 | Inherited |
| $K^{\mathrm{starch}}_{i}$ | 10.00 | Inherited |
| $K^{\mathrm{starch}}_{i,\mathrm{3PGA}}$ | 0.10 | Inherited |
| $K^{\mathrm{starch}}_{i,\mathrm{F6P}}$ | 0.02 | Inherited |
| $K^{\mathrm{starch}}_{i,\mathrm{FBP}}$ | 0.02 | Inherited |
| $K^{\mathrm{FBPase}}_{i,\mathrm{F6P}}$ | 0.70 | Inherited |
| $K^{\mathrm{FBPase}}_{i,\mathrm{P_i}}$ | 12.00 | Inherited |
| $K^{\mathrm{PRK}}_{i,\mathrm{3PGA}}$ | 2.00 | Inherited |
| $K^{\mathrm{PRK}}_{i,4}$ | 2.50 | Inherited |
| $K^{\mathrm{PRK}}_{i,5}$ | 0.40 | Inherited |
| $K^{\mathrm{PRK}}_{i,\mathrm{P_i}}$ | 4.00 | Inherited |
| $K^{\mathrm{PRK}}_{i,\mathrm{RuBP}}$ | 0.70 | Inherited |
| $K^{\mathrm{Rubisco}}_{i,\mathrm{3PGA}}$ | 0.04 | Inherited |
| $K^{\mathrm{Rubisco}}_{i,\mathrm{FBP}}$ | 0.04 | Inherited |
| $K^{\mathrm{Rubisco}}_{i,\mathrm{NADPH}}$ | 0.07 | Inherited |
| $K^{\mathrm{Rubisco}}_{i,\mathrm{P_i}}$ | 0.90 | Inherited |
| $K^{\mathrm{Rubisco}}_{i,\mathrm{SBP}}$ | 0.075 | Inherited |
| $K^{\mathrm{SBPase}}_{i,\mathrm{P_i}}$ | 12.00 | Inherited |
| $K^{\mathrm{export}}_{m,\mathrm{DHAP}}$ | 0.077 | Inherited |
| $K^{\mathrm{starch}}_{m,\mathrm{ATP}}$ | 0.08 | Inherited |

*Continued on next page*

Table S4: All model parameters (continued)

| Parameter | Value | Parameter category |
|---|---|---|
| $K^{\text{starch}}_{m,\text{G1P}}$ | 0.08 | Inherited |
| $K^{\text{export}}_{m,\text{GAP}}$ | 0.075 | Inherited |
| $K^{\text{export}}_{m,\text{3PGA}}$ | 0.25 | Inherited |
| $K^{\text{FBPase}}_{m,\text{S}}$ | 0.03 | Inherited |
| $K^{\text{TPT}}_{m,\text{P}_\text{i}}$ | 0.63 | Inherited |
| $K^{\text{TPT}}_{m,\text{P}_\text{i,ext}}$ | 0.74 | Inherited |
| $K^{\text{PRK}}_{m,\text{ATP}}$ | 0.05 | Inherited |
| $K^{\text{PRK}}_{m,\text{Ru5P}}$ | 0.05 | Inherited |
| $K^{\text{Rubisco}}_{m,\text{CO}_2}$ | 0.0107 | Inherited |
| $K^{\text{Rubisco}}_{m,\text{RuBP}}$ | 0.02 | Inherited |
| $K^{\text{SBPase}}_{m,\text{S}}$ | 0.0130 | Inherited |
| $k^{\text{re}}_{\text{ALD,SBP}}$ | $8.00 \times 10^8$ | Inherited |
| $k^{\text{re}}_{\text{ALD,FBP}}$ | $8.00 \times 10^8$ | Inherited |
| $k^{\text{re}}_{\text{G6PI}}$ | $8.00 \times 10^8$ | Inherited |
| $k^{\text{re}}_{\text{GAPDH}}$ | $8.00 \times 10^8$ | Inherited |
| $k^{\text{re}}_{\text{PGM}}$ | $8.00 \times 10^8$ | Inherited |
| $k^{\text{re}}_{\text{PGK}}$ | $8.00 \times 10^8$ | Inherited |
| $k^{\text{re}}_{\text{RPI}}$ | $8.00 \times 10^8$ | Inherited |
| $k^{\text{re}}_{\text{RPE}}$ | $8.00 \times 10^8$ | Inherited |
| $k^{\text{re}}_{\text{TK,F6P}}$ | $8.00 \times 10^8$ | Inherited |
| $k^{\text{re}}_{\text{TK,S7P}}$ | $8.00 \times 10^8$ | Inherited |
| $k^{\text{re}}_{\text{TPI}}$ | $8.00 \times 10^8$ | Inherited |
| $[\text{P}_\text{i}]_\text{ext}$ | 0.50 | Inherited |
| $\text{P}_\text{i,tot}$ | 17.05 | Inherited |
| **Ascorbate–glutathione cycle and sinks** | | |
| $\text{ASC}_\text{tot}$ | 10.00 | Inherited |
| $E_{0,\text{DHAR}}$ | 0.0017 | Inherited |
| $E_{0,\text{GR}}$ | 0.0014 | Inherited |
| $E_{0,\text{MDAR2}}$ | 0.0020 | Inherited |
| $\text{GSH}_\text{tot}$ | 10.00 | Inherited |
| $K_\text{DHAR}$ | 0.50 | Inherited |
| $k^{\text{cat}}_{\text{DHAR}}$ | 142.00 | Inherited |
| $k^{\text{cat}}_{\text{GR}}$ | 595.00 | Inherited |
| $k^{\text{cat}}_{\text{MDAR2}}$ | 300.00 | Inherited |
| $k^{\text{f}}_1$ | $1.00 \times 10^4$ | Inherited |
| $k^{\text{f}}_2$ | $1.00 \times 10^4$ | Inherited |
| $k^{\text{f}}_3$ | 2510.00 | Inherited |
| $k^{\text{f}}_4$ | $1.00 \times 10^4$ | Inherited |

*Continued on next page*

Table S4: All model parameters (continued)

| Parameter | Value | Parameter category |
|---|---:|---|
| $k_5^{\mathrm{f}}$ | 2510.00 | Inherited |
| $k_{\mathrm{MDAR1}}$ | 500.00 | Inherited |
| $K_{m,\mathrm{DHA}}^{\mathrm{DHAR}}$ | 0.07 | Inherited |
| $K_{m,\mathrm{GSH}}^{\mathrm{DHAR}}$ | 2.50 | Inherited |
| $K_{m,\mathrm{GSSG}}^{\mathrm{GR}}$ | 0.20 | Inherited |
| $K_{m,\mathrm{NADPH}}^{\mathrm{GR}}$ | 0.0030 | Inherited |
| $K_{m,\mathrm{MDA}}^{\mathrm{MDAR2}}$ | 0.0014 | Inherited |
| $K_{m,\mathrm{NADPH}}^{\mathrm{MDAR2}}$ | 0.023 | Inherited |
| $k_1^{\mathrm{r}}$ | 220.00 | Inherited |
| $k_2^{\mathrm{r}}$ | 4000.00 | Inherited |
| $k_4^{\mathrm{r}}$ | 4000.00 | Inherited |
| $X_{\mathrm{T}}$ | 0.07 | Inherited |

#### 1.5 initial conditions

Table S5: Initial conditions of the model variables

| Variable | Symbol | Initial value |
|---|---|---|
| **Calvin–Benson–Bassham cycle** | | |
| 3-Phosphoglycerate (3PGA) | [3PGA] | 0.92 |
| 1,3-Bisphosphoglycerate (BPGA) | [BPGA] | 0.00038 |
| Glyceraldehyde 3-phosphate (GAP) | [GAP] | 0.0058 |
| Dihydroxyacetone phosphate (DHAP) | [DHAP] | 0.13 |
| Fructose 1,6-bisphosphate (FBP) | [FBP] | 0.0053 |
| Fructose 6-phosphate (F6P) | [F6P] | 0.29 |
| Glucose 6-phosphate (G6P) | [G6P] | 0.66 |
| Glucose 1-phosphate (G1P) | [G1P] | 0.0384 |
| Sedoheptulose 1,7-bisphosphate (SBP) | [SBP] | 0.0111 |
| Sedoheptulose 7-phosphate (S7P) | [S7P] | 0.15 |
| Erythrose 4-phosphate (E4P) | [E4P] | 0.0067 |
| Xylulose 5-phosphate (X5P) | [X5P] | 0.021 |
| Ribose 5-phosphate (R5P) | [R5P] | 0.0352 |
| Ribulose 1,5-bisphosphate (RuBP) | [RuBP] | 0.11 |
| Ribulose 5-phosphate (Ru5P) | [Ru5P] | 0.0141 |
| **Energy and redox carriers** | | |
| ATP | [ATP] | 1.46 |
| NADPH | [NADPH] | 0.56 |
| Ferredoxin, oxidised | $[Fd_{ox}]$ | 3.72 |
| Plastocyanin, oxidised | $[PC_{ox}]$ | 1.81 |
| Plastoquinone, oxidised | $[PQ_{ox}]$ | 10.30 |
| **Photosystems** | | |
| PSII state $B_0$ | $[B_0]$ | 1.94 |
| PSII state $B_1$ | $[B_1]$ | $9.79 \times 10^{-8}$ |
| PSII state $B_2$ | $[B_2]$ | 0.56 |
| PSI: P700 $F_A$ | $[P700\,F_A]$ | 1.51 |
| PSI: $P700^+$ $F_A^-$ | $[P700^+ F_A^-]$ | 0.0192 |
| PSI: P700 $F_A^-$ | $[P700\,F_A^-]$ | 0.0281 |
| **Non-photochemical quenching** | | |
| Light-harvesting complex | [LHC] | 0.78 |
| PsbS, deprotonated | $[PsbS_{deprot}]$ | 0.97 |
| Violaxanthin | [Vx] | 0.96 |
| **Reactive oxygen species and ascorbate–glutathione cycle** | | |

*Continued on next page*

Table S5: Initial conditions of the model variables (continued)

| Variable | Symbol | Initial value |
|---|---|---|
| Hydrogen peroxide | $[H_2O_2]$ | $1.2 \times 10^{-7}$ |
| Monodehydroascorbate (MDA) | [MDA] | $2.04 \times 10^{-7}$ |
| Dehydroascorbate (DHA) | [DHA] | $1.03 \times 10^{-11}$ |
| Glutathione disulfide (GSSG) | [GSSG] | $5 \times 10^{-12}$ |
| **Regulation** | | |
| Thioredoxin, oxidised | $[\mathrm{Trx_{ox}}]$ | 0.93 |
| CBB enzymes, inactive | $[\mathrm{E_{inactive}}]$ | 5.50 |
| ATP synthase, active fraction | $\mathrm{a_{ATPsynth}}$ | 0 |
| **pH, membrane potential and ions** | | |
| Luminal pH | $\mathrm{pH_{lumen}}$ | 7.00 |
| Stromal pH | $\mathrm{pH_{stroma}}$ | 7.00 |
| Membrane potential (V) | $\Delta\psi$ | 0 |
| Stromal potassium | $[\mathrm{K^+_{stroma}}]$ | 34.50 |
| Luminal potassium | $[\mathrm{K^+_{lumen}}]$ | 34.50 |
| Stromal chloride | $[\mathrm{Cl^-_{stroma}}]$ | 15.65 |
| Luminal chloride | $[\mathrm{Cl^-_{lumen}}]$ | 15.65 |
